# Variation in *Melitaea cinxia* gut microbiota is phylogenetically highly structured but only mildly driven by host plant microbiota, sex or parasitism

**DOI:** 10.1101/510446

**Authors:** G. Minard, G Tikhonov, O. Ovaskainen, M. Saastamoinen

**Affiliations:** Organismal and Evolutionary Biology Research Programme, University of Helsinki, Helsinki, Finland; Université de Lyon, Lyon, France; Ecologie microbienne, UMR CNRS 5557, UMR INRA 1418, VetAgro Sup, Université Lyon 1, Villeurbanne, France; Centre for Biodiversity Dynamics, Department of Biology, Norwegian University of Science and Technology. N-7491 Trondheim, Norway

**Keywords:** Microbiome, metabolome, insect, arthropod, microbial ecology, community modeling, joint species distribution model

## Abstract

**Originality-Significance Statement:** The factors contributing to the assembly of microbiota in animals are extremely complex, and thus a comprehensive understanding of the mechanisms shaping host-associated microbial communities in natural ecosystems requires extensive ecological studies and appropriate statistical methods. In this study, we investigated the bacterial microbiota associated with the caterpillars of the Glanville fritillary (*Melitaea cinxia*), which is a long-term-studied ecological model system. We assessed the structure of variation in both occurrence and abundance of gut microbial communities of individuals collected in the wild with joint-species modelling, with the aim to relate the microbial community structure with multiple potentially impacting covariates: host plant microbiota and metabolites, hosts’ sex, potential parasitoid infection, and family structure. These covariates exhibited substantial correlation with multiple microbial taxa’s occurrences, which correlations were consistent for phylogenetically related groups of taxa, but varied across the whole microbial community; on the contrary, only few correlations were found with taxa’s abundances. The dominating co-occurrence pattern of microbiota assembly, which effectively split caterpillar individuals into two distinct groups, was, however, unrelated to any of the considered covariates.

**Summary:** Understanding of what ecological factors shape intraspecific variation of insect microbiota is still relatively poor. In Lepidopteran caterpillars, microbiota is assumed to be mainly composed of transient bacterial symbionts acquired from the host plant. We sampled Glanville fritillary (*Melitaea cinxia*) caterpillars from natural populations to describe the microbiome and to identify potential factors that determine the structure of the microbial community, including the sex of the host, the impact of parasitoid infection, and the possible link between host plant and caterpillar microbiota. Our results demonstrate high variability of microbiota composition even among caterpillars that shared the same host plant individual. The observed variation in microbiota composition is partially attributed to the measured properties of the host or its plant microbial and chemical composition, and is aligned with microbial phylogenetic structure, with related taxa exhibiting similar patterns. However, the prevailing part of the observed variation was not associated with any of the assessed characteristics, although it followed a pronounced segregation structure: in some caterpillars the microbial communities were dominated by several related Enterobacteriaceae taxa, while in others these taxa were absent. Our results challenge previous findings that the host plant properties are the major drivers of microbiota communities of insect herbivores.

## Introduction

All animals interact with microorganisms (McFall-Ngai *et al.,* 2013), with interactions between hosts and their microbes ranging from mutualistic to competitive (Douglas, 2010). Insects harbor highly diversified host-symbiont interactions with various examples of fitness benefits (Douglas, 2011), such as the control of the host’s reproduction (Werren *et al.,* 2008; Engelstädter and Hurst, 2009), the enhancement of nutrition via effects on the digestion process (Warnecke *et al.,* 2007), the degrading of toxic metabolites (Kikuchi *et al.,* 2012; Ceja-Navarro *et al.,* 2015), and the production of nutrients essential for the host (Akman Gunduz and Douglas, 2009; Salem *et al.,* 2014). Endosymbionts can also protect their hosts against abiotic stressors and pathogens (Montllor *et al.,* 2002; Dunbar *et al.,* 2007; King *et al.,* 2016). The literature may, however, be biased towards mutualistic and parasitic/pathogenic interactions, whereas commensal or neutral interactions may be understudied or underreported. In general, the microbiota is a multilayer system in which prevalent members compose the core microbiota and a more flexible pool of microbial members compose the non-core community (Shapira, 2016).

Host-microbiota interactions are often complex, involve multiple taxa and multiple transmission processes, and consequently laboratory-based studies may fail to realistically portray natural systems. Indeed, several studies have highlighted pronounced differences in the microbiota of laboratory-reared versus field-captured individuals (Rani *et al.,* 2009; Staubach *et al.,* 2013; Tinker and Ottesen, 2016). Characterizing and determining the impact of microbiota in natural populations remains challenging, due to the multiple confounding factors that can affect the microbiota composition. Consequently, we still know little of the ecological factors that shape among-individual variation of microbial communities in natural populations. Another challenge is related to data analyses: microbiota data typically include large numbers of taxonomical units, most of which are rare, complicating the use of conventional statistical frameworks.

The gut microbiota of insects is often highly heterogenic both among and within species, with relatively high variation reported even across different gut sections (Douglas, 2015). The consumed diet has been suggested to be the major determinant of the microbiota composition, as it can shape the microbial communities both directly (e.g. acquisition of food-associated microorganisms or growth of microorganisms that utilize the consumed food) and indirectly (e.g. through impacts on immunity, anatomy or digestive function; Douglas, 2015). However, several studies that controlled for the transient effects of diet (e.g. in fruit flies and Asian tiger mosquitoes), still reported strong inter-individual variation in the microbiota composition (Minard *et al.,* 2015; Adair *et al.,* 2018), suggesting the importance of diet-unrelated factors. For example, gut microbiota can be acquired via maternal or horizontal transmission (Engel and Moran, 2013), influenced by host genotype or environmental conditions unrelated to food (Yun *et al.,* 2014), or they can be driven mainly by stochastic processes (Douglas, 2015; Zeng *et al.,* 2015). In Lepidoptera, there is only little evidence on the transfer of symbiotic bacteria among individuals (Paniagua Voirol *et al.,* 2018). Consistently, the Lepidopteran gut microbiome has been shown to be highly variable compared to other insect orders, with only few resident bacteria (Hammer *et al.,* 2017). The impact of the gut microbiota on the life history traits of Lepidoptera has been questioned, even though the general knowledge on the bacterial associations across species is still very limited (see Paniagua Voirol *et al.,* 2018 for a review).

To improve our understanding of the potential ecological determinants influencing associations between insect hosts and their gut symbionts, we exploit here the natural metapopulation of the Glanville fritillary butterfly (*Melitaea cinxia)* in the Åland islands, Finland. With *M. cinxia* caterpillars and their *Plantago lanceolata* host plants sampled across this system at a single timepoint, our overall aim is to associate the midgut microbiota of the caterpillars with ecological variables and thus to identify potential drivers of variation that could impact these communities. In particular, we ask (1) whether there is a correspondence between the host plant microbiota and that of the caterpillar microbiota; (2) whether the host plant microbiota and the caterpillar microbiota are influenced by the metabolite profile of the host plant; (3) whether the caterpillar microbial communities are structured according to the sex and parasitoid infection status of the host; (4) whether after accounting for the above mentioned factors, variation in these communities is structured according to caterpillar families or is idiosyncratic among individuals independent of the family structure; and (5) whether the variation in the microbiota with respect to the questions 1-4 is phylogenetically structured. Further, to examine if and how microbial variation influences the fitness of the host, we ask (6) whether the over-winter survival of caterpillar nests can be explained by microbiota composition. To address these questions, we apply a joint species distribution model that allows us to evaluate both species- and community-level responses to the abovementioned covariates, as well as co-occurrence patterns of the microbiota both at the levels of individual caterpillars and caterpillar families.

## Results

### Factors influencing caterpillar microbiota

Overall, the caterpillar microbiota was composed of variable microbiota among which the dominant taxa (>1% of the relative abundance across all samples) were *Uruburella, Cloacibacterium, Moraxella, Acinetobacter, Dermacoccus, Hymenobacter, Corynebacterium, Paracoccus, Wolbachia, Methylobacterium,* and unclassified Actinobacteria, Enterobacteriaceae and Corynebacteriaceae (Fig. 1A, Fig. 2). *Uruburuella* was most prevalent but still detected in only 58.8% of the samples, suggesting no core microbiota across all individuals. To investigate the potential ecological factors explaining variation in the occurrences and abundances of microbial taxa, we used joint-species modelling framework.

**Figure 1.**
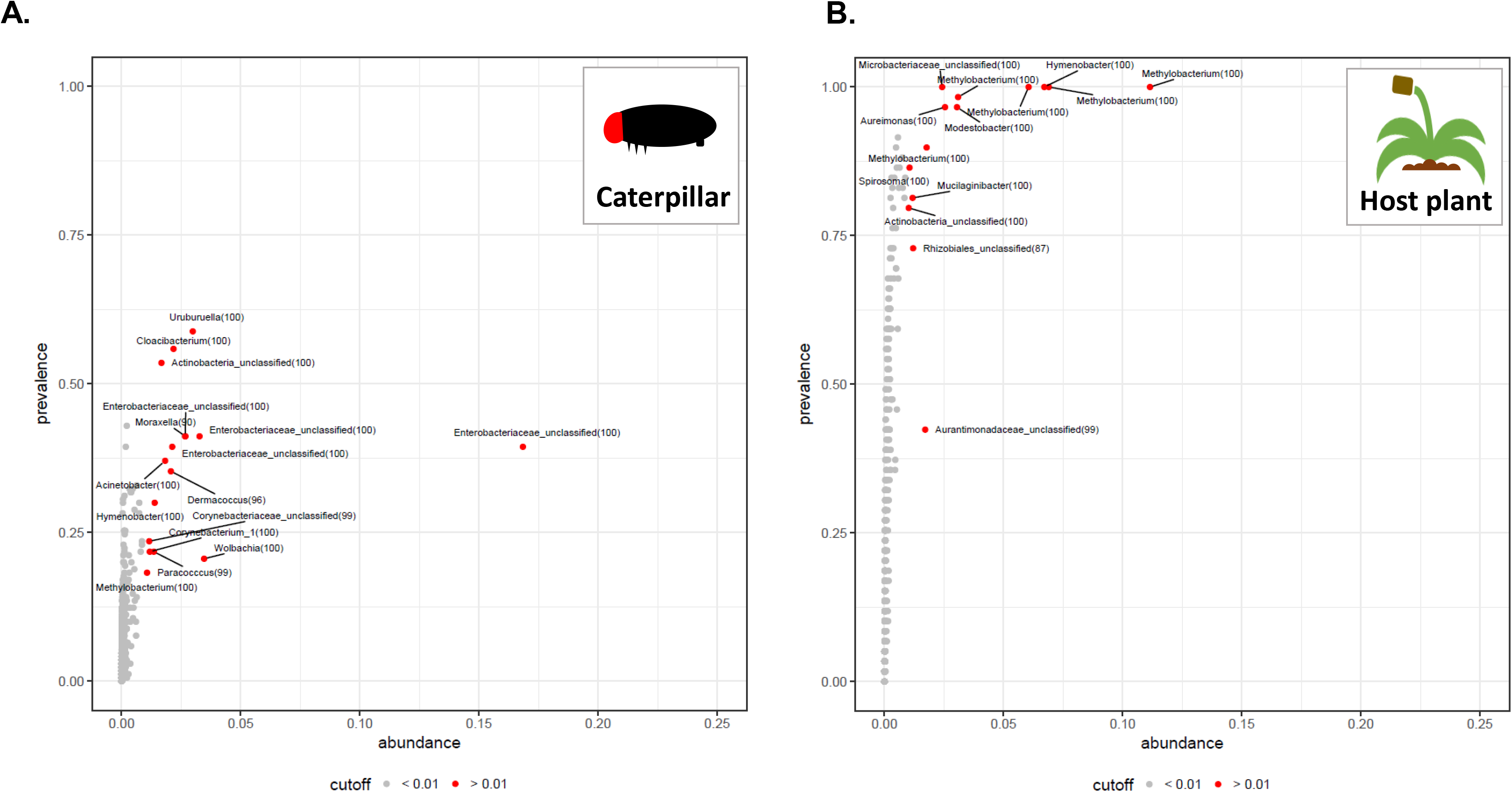
Prevalence and average abundance of the bacteria within caterpillar midguts (A) and plant leaves (B). The OTUs (dots) are represented according to the proportion of individual samples in which they were detected (prevalence) and their average relative abundance across all the samples (abundance). The classification of the most abundant OTUs (cutoff > 0.1) is provided. The bootstrap associated with each taxonomical classification is reported under brackets.

**Figure 2.**
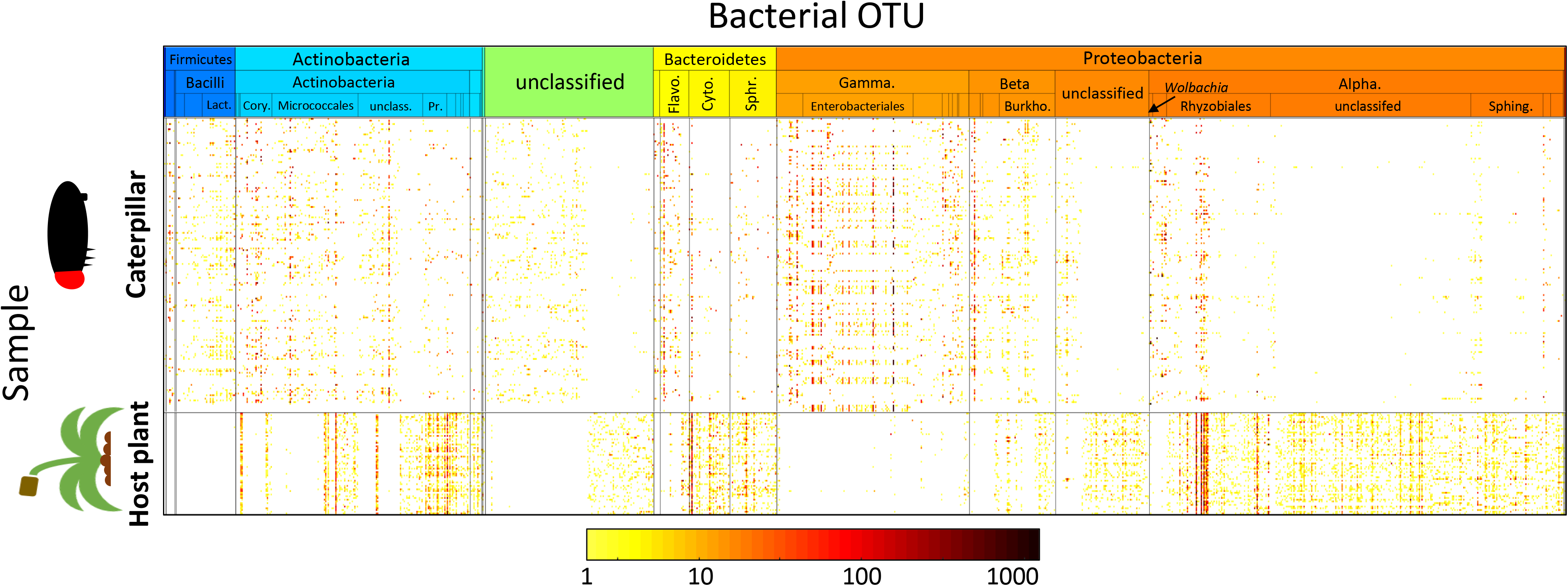
Abundances of bacterial OTUs in caterpillar and plant samples. The OTUs (columns) have been ordered according to their taxonomical classification (for details, see Supplementary Table S2). The color scale shows OTU abundance (number of normalized sequences) for each caterpillar and plant sample on a logarithmic scale, and white color indicates absence of OTU in given sample.

Averaged over the whole community, the presence-absence part of the caterpillar model had only little predictive power through its fixed effects (Prediction P1; Supplementary Table S1). In the line with this, caterpillar sex, parasitoid infection or the host plant’s bacterial or metabolic composition did not show a community-consistent correlation with the presence-absence patterns of the bacterial community ([5%,95%] credibility interval for community-level mean value of species response overlapped with zero, Table 1). Accounting for the residual species-to-species associations substantially increased the predictive power, which however remained highly variable over the OTUs (Prediction P2; Supplementary Table S1), but was only slightly worse than the explanatory power of the full model (Prediction P3; Supplementary Table S1). These results suggest that the modelled associations among bacterial taxa (OTUs) represent a true biological signal instead of merely model overfitting artefact. In the abundance model, both the fixed effects and the species-to-species associations contributed roughly equally (Supplementary Table S1). However, similarly to the presence-absence model, none of the fixed effects had a consistent correlation with the abundance pattern, and thus the influences of the environmental covariates were taxon-specific. The variance partitioning of additive Gaussian components in HMSC’s latent predictor mirrored the results of predicted power comparison (Fig. 3).

**Table 1.**
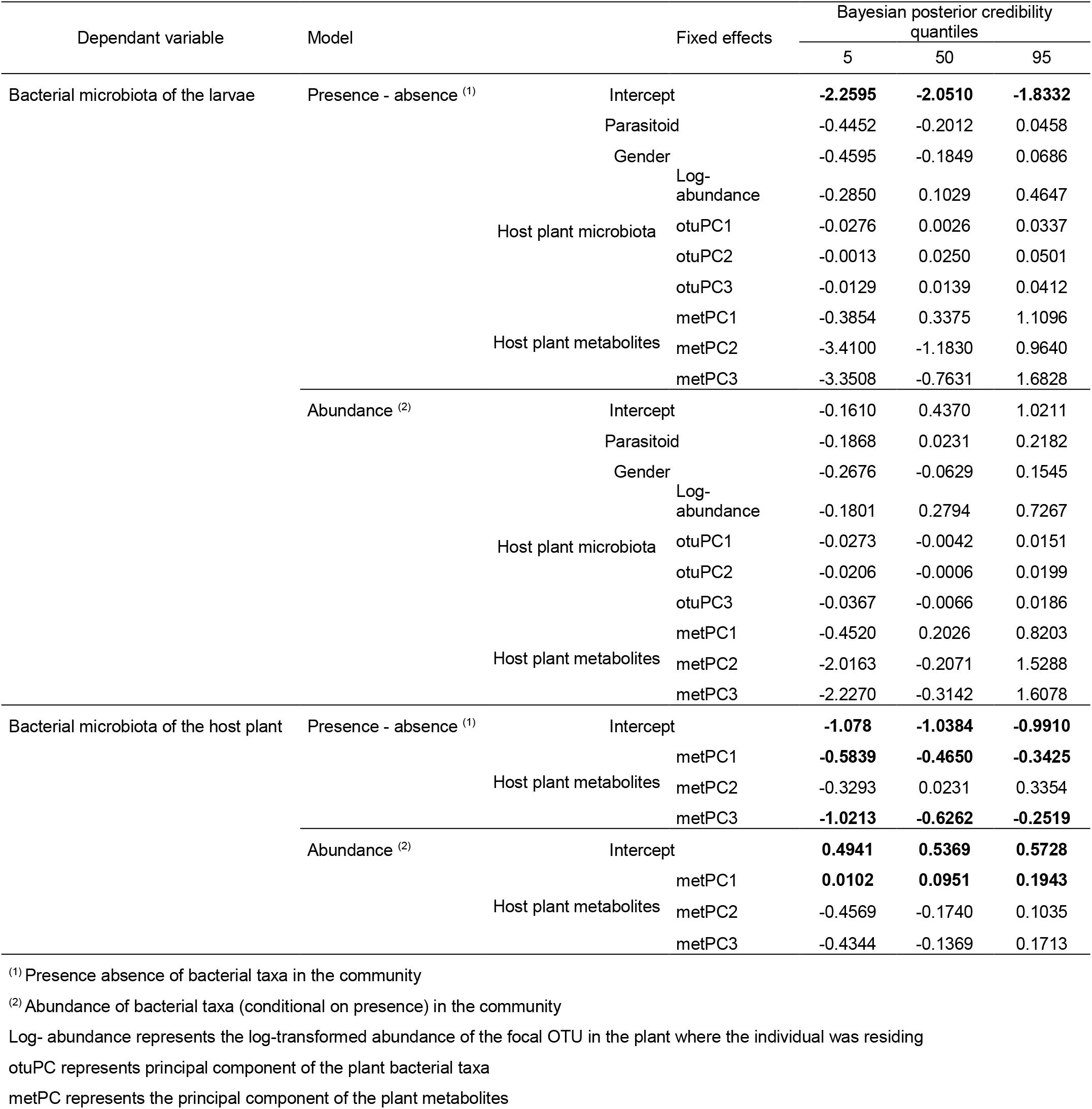
Estimates of the community-consistent fixed effects included in the modeling of the bacterial communities associated with larvae and their host plant.

**Figure 3.**
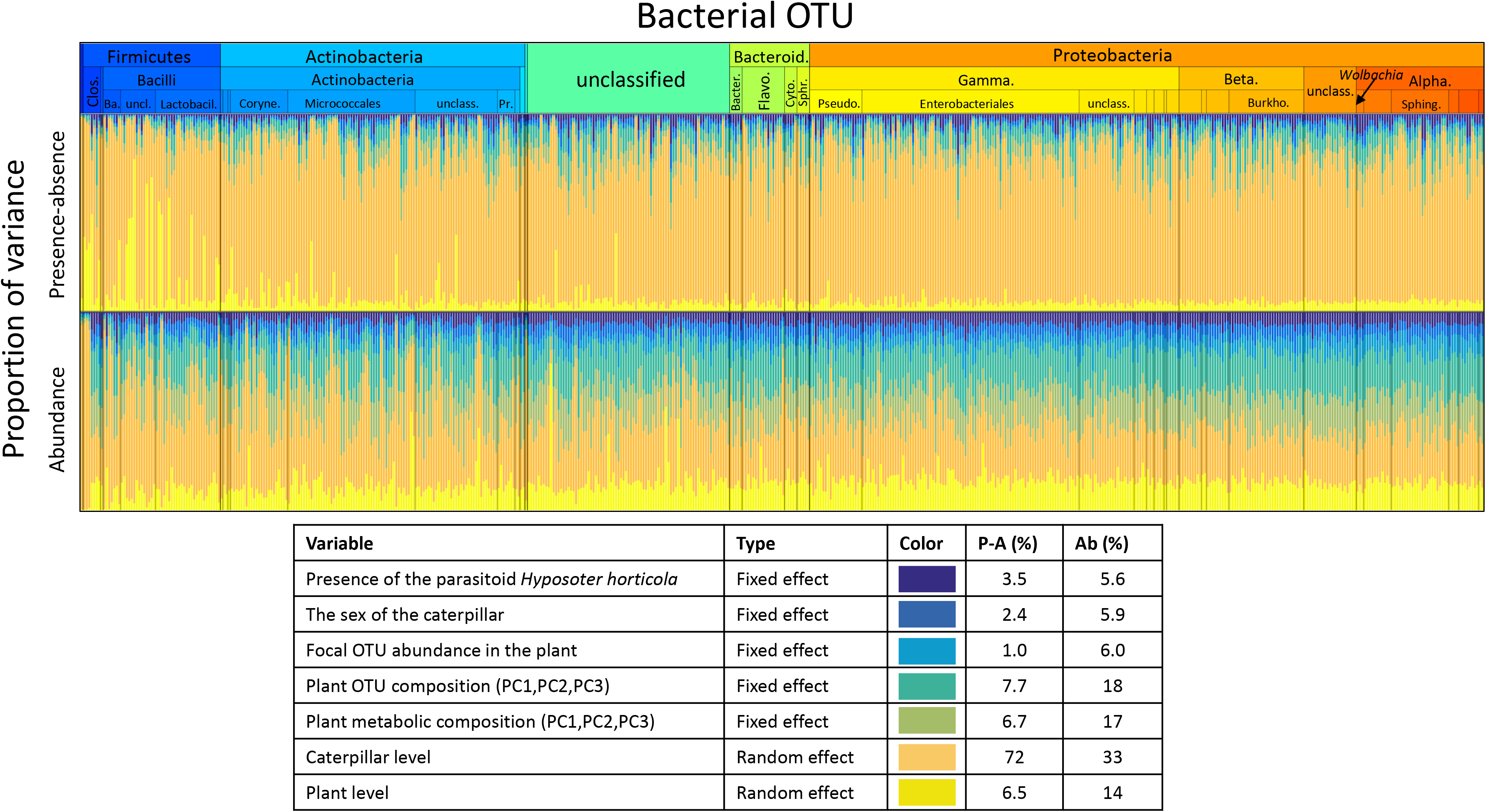
Partitioning of variation in caterpillar microbiota to components explained by different types of fixed and random effects. The colored bars show, for each OTU, the proportions of variance attributed to each group of explanatory variables. The average variance proportions over OTUs are shown in the legend, with P-A corresponding to the presence-absence and Ab to the abundance model. The order of OTUs is following the ordering of Fig. 2 except for OTUs that were recorded only in plant samples and are omitted here (for details, see Supplementary Table S2). See *Statistical Methods* for a full description of the included fixed and random effects.

Despite the generally low proportion of explained variance (Fig. 3), a substantial proportion of OTUs showed positive or negative responses to the fixed effects (Fig. 4). Specifically, the occurrence probabilities of many OTUs decreased with the presence of the parasitoid infection (mostly Clostridia, Alphaproteobacteria and Betaproteobacteria), and were lower in males than in females (mostly Rhodobacteriales and Neisseriales). The presence of *Wolbachia,* on the other hand, was positively associated with the parasitoid infection of the caterpillars. Only a minority of the OTUs classified as *Hymenobacter* and *Methylobacterium* showed increased occurrence probability with the increased abundance of the focal OTU in the host plant. Unlike in presence-absence model, in the abundance model only a small proportion of the individual OTUs’ responses gained substantial statisticalsupport (Fig. 4).

**Figure 4.**
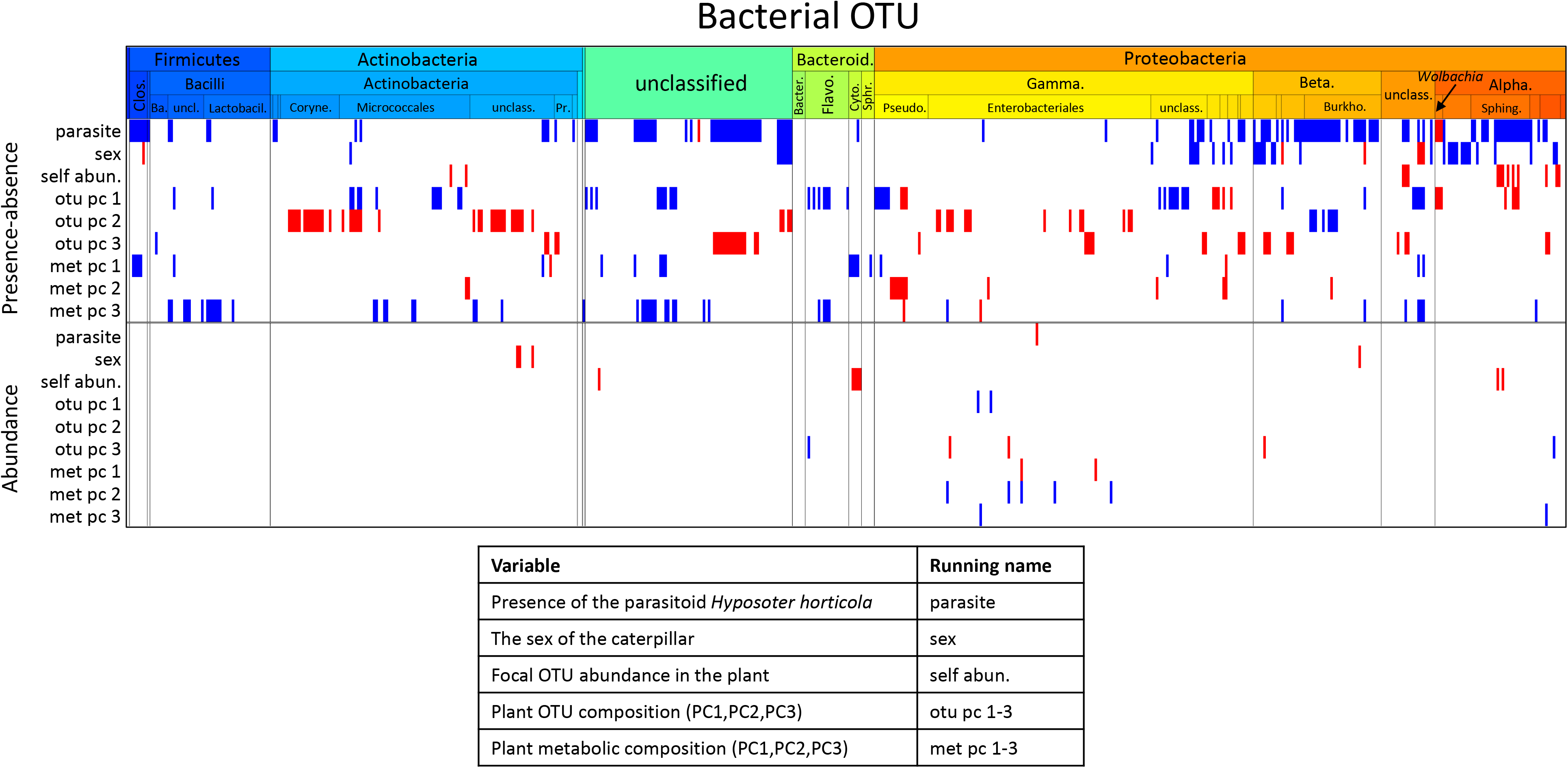
Influence of measured covariates on caterpillar microbiota. Regression coefficients that were estimated to be positive (respectively, negative) with 95% credibility level are shown by red (respectively, blue). The ordering of OTUs is identical to that of Fig. 2 except for OTUs that were recorded only in plant samples and are omitted here (for details, see Supplementary Table S2). The covariates included in the model are listed in the legend alongside with their running names used in axis labelling.

Phylogenetically related bacterial taxa were estimated to have similar responses to the covariates: the strength of the phylogenetic signal was 0.98±0.002 (posterior mean ± posterior standard deviation) in the presence-absence model and 0.86±0.025 in the abundance model. This shows that closely related bacteria had similar niches in the sense that they responded similarly to the fixed effects included in the model. This result is clearly visible in Fig. 4, that represents the responses of bacterial taxa ordered by taxonomy, and where the positive and negative effects (the red and blue colors) are presented as contiguous blocks rather than randomly distributed across the OTUs. The majority of the Betaproteobacteria, for example, have lower occurrence probability when the individual is infected by the parasitoid. The occurrence of the microbial OTUs were phylogenetically structured not only with respect to the measured covariates, but also in their residual variation, as the OTUs split into two groups in a markedly pronounced manner (Fig. 5A). One of these two groups consisted, with minor exceptions, of the Enterobacteriaceae family, and the other group of the remaining taxa including *Uruburella, Cloacibacterium, Moraxella, Acinetobacter, Dermacoccus, Hymenobacter, Corynebacterium, Paracoccus.* Thus, some of the caterpillars were characterized by a high representation of Enterobacteriaceae in their microbiota, while the remaining individuals were characterized by a low representation of Enterobacteriaceae. Given its dominant role in variance partitioning, this pattern is the strongest signal related to OTU occurrences in our data (Fig. 5A), and its validity is supported by similar results of a complementary analysis based on Dirichlet mixture modelling (Supporting Information, Fig. S1AC). In contrast to the strong patterns recorded in the presence-absence model, only few statistically supported associations were found in the abundance model (Fig. 5B). Quite unexpectedly, the caterpillar family structure was estimated to play only a minor role on the microbial community structure, as both for the presence-absence and abundance models it was attributed with minor proportion of variance (Fig. 3) and almost no statistically supported associations were found.

**Figure 5.**
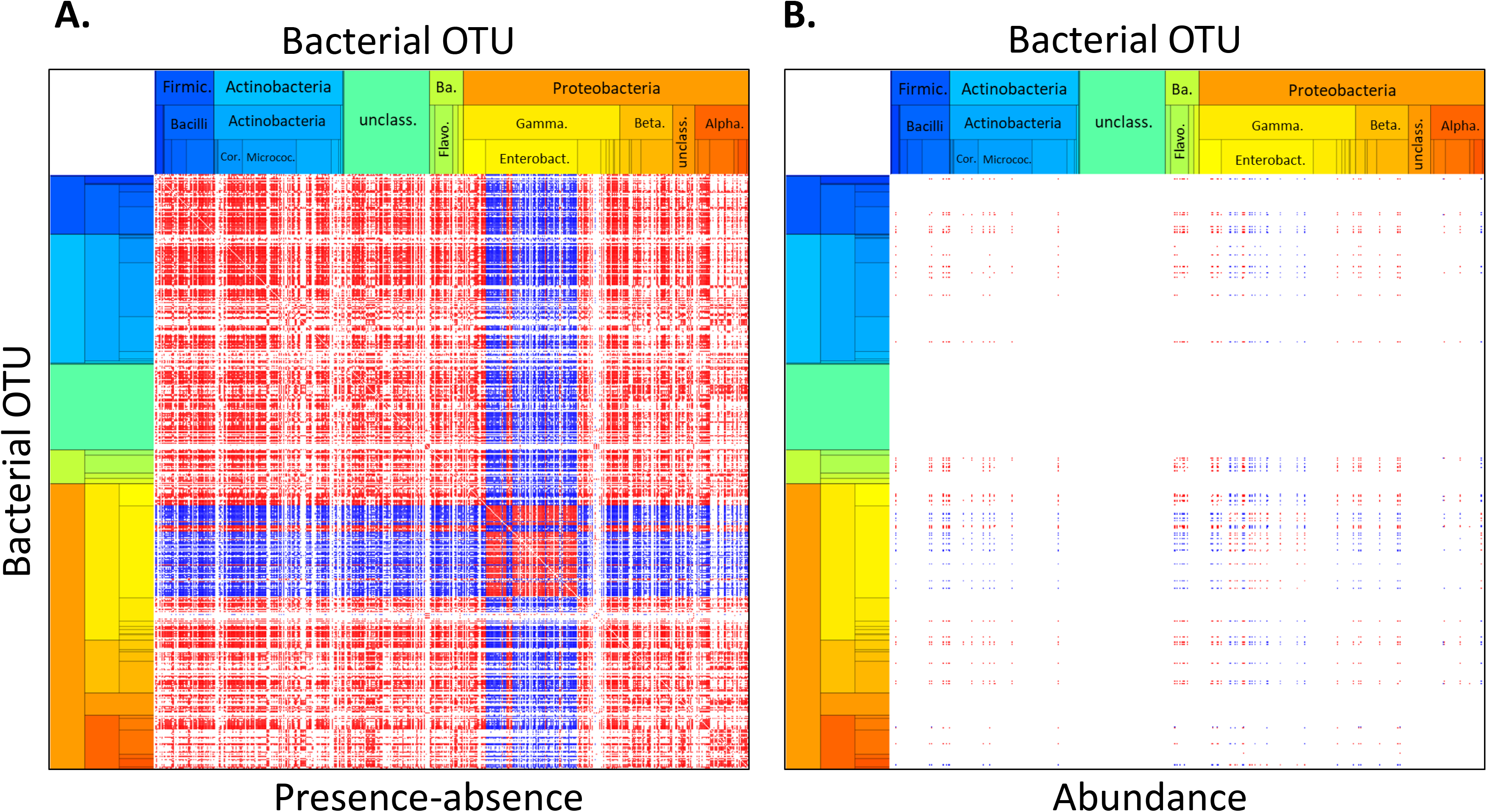
Residual associations among caterpillar microbiota. The panels illustrate the caterpillar-level random effects for the presence-absence (A) and abundance (B) parts of the caterpillar model. OTU-pairs for which the residual correlation was estimated to be positive (respectively, negative) with 95% credibility level are shown by red (respectively, blue) color. The ordering of OTUs is identical to that of Fig. 2 except for OTUs that were recorded only in plant samples and are omitted here (for details, see Supplementary Table S2).

To summarize, the variation of bacterial community exhibited a complex structure, which had both inter-individual and inter-OTU patterns, and did not support a simple characterization. In terms of variation among caterpillar individuals, we found that neither the fixed effects assessed nor the family relationships were capable to explain the very strong segregation of individuals into two groups with very distinct microbial composition: about 40% of the individuals were characterized by the microbiota with a co-occurrence of phylogenetically related Enterobacteriaceae, whereas the rest of the individuals were characterized by a more complex microbial community, composed of *Uruburella, Cloacibacterium, Moraxella, Acinetobacter, Dermacoccus, Hymenobacter, Corynebacterium, Paracoccus, Wolbachia, Methylobacterium,* and some unclassified Actinobacteria and Corynebacteriaceae. Although, the fixed effects included in our model accounted for minor part of this variation, and the microbial OTU responses to the fixed effects were not synchronized across whole community, we found that phylogenetically similar OTUs responded to these effects in similar manner.

### Factors influencing host plant foliar microbiota

Contrary to the microbiota within the caterpillar gut, the plant microbiota was composed of highly prevalent bacterial taxa (detected in more than 90% of the collected samples; Fig. 1B). The bacteria in this core microbiota were assigned to *Methylobacterium, Hymenobacter, Aureimonas, Modestobacter,* and an unclassified Microbacteriaceae. Whenever possible, we also assessed the plant metabolome by ^1^H-NMR spectrometry (Fig. 6). The identified metabolites included amino acids (Valine, Threonine, Alanine, Arginine, Glutamate, Glutamine), sugars (Xylose, α-glucose, β-glucose, Sucrose), organic acids (Fumaric acid, acetic acid, cis-aconic acid), ethanol and defensive metabolites (Aucubin, Catalpol and Verbasosides). The latter included both terpenoids (Aucubin, Catalpol) and phenolic compounds (Verbascosides), which constitute the main chemical defense of *Plantago lanceolata* against herbivores and pathogenic microorganisms. Most of the variation in the plant metabolites across samples were explained by unannotated metabolite signals with a chemical shift specific of carbohydrates and amino acid residues (PC1 in Fig. S3) other annotated metabolites including defensive metabolites showed only limited variations (PC1, PC2, PC3 in Fig. S3).

**Figure 6.**
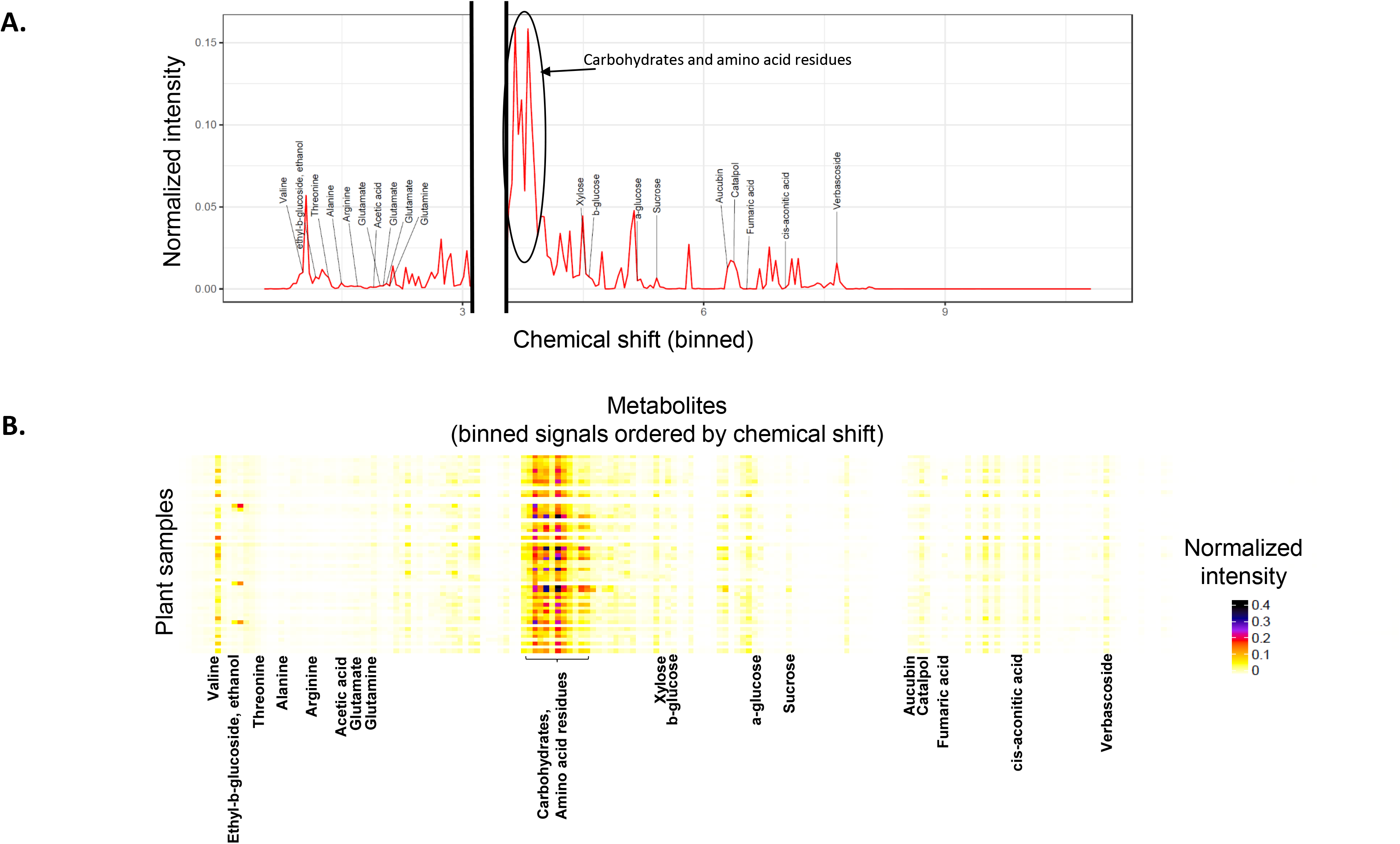
Metabolomic analysis of *Plantagoa lanceolata* leaves. (A) An example spectrum (binned at 0.04 ppm) obtained for a leaf sample of *Pl. lanceolata* and showing the chemical shift of the different annotated metabolites. The gap within the graph represents the removal of the solvent pick. (B) A heatmap of the different binned signals showing their normalized intensity accross all plant samples.

Similar to the OTUs in caterpillars, we applied joint species modelling to assess the presence-absence and the abundance patterns of the bacterial taxa retrieved from the host plant leaves. The plant model showed lower predictive power than caterpillar model, suggesting that the variation in microbial communities was less predictable at plant foliar than in caterpillar guts (Supplementary Table S1). Of the explained variation, the metabolite composition of the plants was the key determinant in both the presence-absence and the abundance models (Table 1, Supplementary Table S1, Supplementary Fig. S4). Here as well, we found a strong phylogenetic structure in explaining how the OTUs responded to fixed effects: the strength of the phylogenetic signal was 0.97±0.01 for presence-absence model and 0.51±0.08 for the abundance model (see Supplementary material, Supplementary Fig. S5, S6). This means that phylogenetically related bacteria responded in a similar manner to the variation in the plant metabolite composition. In particular, lower residues of carbohydrates and amino acids (PC1 in Fig. S3) were negatively associated with Alphaproteobacteria and Actinobacteria (PC1 of the presence absence model in Fig. S7). Contrary to the microbial communities in the caterpillar, unexplained associations between bacteria OTUs were much weaker. The Dirichlet-multinomial modelling results for plant-inhabiting communities indicated that this variation is best explained by a single-component distribution (Supporting Information, Fig. S1B).

### Influence of microbiota on overwinter survival

As the caterpillars of the Glanville fritillary overwinter gregariously, mainly in family groups, we were interested in testing whether the microbiota composition of the samples collected from the field would correlate with the survival of the siblings remaining in the wild. This would have demonstrated an important fitness benefit of the microbiota composition in wild populations. However, we did not find the over-wintering mortality of families to be related to the microbiota composition or the metabolite profile of the host plants in which the caterpillars were residing on (see Supplementary Information).

## Discussion

Symbionts that are highly competitive, strongly adapted to their host, and frequently colonize host populations form the core microbiota shared among individuals of the same species (Shapira, 2016). On the contrary, symbionts that are competitively inferior, less adapted to the intestinal conditions (e.g. pH, and digestive enzymes), and/or are rarely acquired or transmitted among individuals, tend to form a pool of transient bacteria that consequently are subject to higher fluctuations among hosts (Shapira, 2016; Macke *et al.,* 2017). We show that the natural midgut microbial community of *M. cinxia* caterpillar is highly variable, and that only a minor proportion of that variation is related to the measured caterpillar’s traits or the properties of the host plant in which it was feeding. In contrast, we documented a strong co-occurrence patterns of OTUs at the level of caterpillar individual, which could not be attributed to any of the covariates included in our analyses. These co-occurrence patterns in the microbiota were strongly phylogenetically structured, suggesting two mutually exclusive groups of bacterial communities. One of these groups consisted of mainly Enterobacteriaceae, whereas the other group consisted of the remaining taxa. Enterobacteriaceae contains several taxa specifically associated with animal digestive system with a broad range of host-microbe interactions ranging from pathogenic to mutualistic (Douglas, 1998; Weiss *et al.,* 2006; Chandler *et al.,* 2011; Parmentier *et al.,* 2016). Enterobacteriaceae are one of the most widespread bacteria also known to be associated with Lepidoptera (Paniagua Voirol *et al.,* 2018), and in *Heliconius erato,* for example, they dominate the gut microbiota already in the early developmental stages (Hammer *et al.,* 2014). Consistent with our results, the microbiota of *Drosophila melanogaster* has also been shown to be phylogenetically structured (Adair *et al.,* 2018). In general, Lepidopteran associated microbiota are suggested to be highly variable (Staudacher *et al.,* 2016): a study on caterpillars representing 124 Lepidopteran species showed high inter- and intraspecific variation in the gut microbiota, with a poorly abundant core microbiota (Hammer *et al.,* 2017). The dominance of co-occurring taxa, such as Enterobacteriaceae in our study, may be driven by several factors, such as priority effects (dominance of a group of microbes that were the first to colonize the gut), the specific association of bacteria involved in mutualistic interactions, or by a niche overlap among the co-occurring bacteria that grow under similar conditions (Kennedy and Bruns, 2005; Peay *et al.,* 2012; Sprockett *et al.,* 2018). Due to the limitation in the biological material, we could not quantify absolute abundance with e.g. qPCR. As our results are based on relative abundances, we cannot exclude the possibility that e.g. the individuals have otherwise a uniform microbiota but some individuals are additionally massively colonized by Enterobacteriaceae.

### Sex and parasitoid infection are correlated with variation of marginal bacterial taxa

The occurrence of OTUs within Rhodobacterales and Neisseriales orders was generally higher in female than male caterpillars. Due to the absence of sexual dimorphism and proper genetic markers, most studies conducted on immature developmental stages of insects fail to consider sex differences in the microbiome. However, sex-specific differences may greatly impact the microbiota from early caterpillar instar onwards. In the silkworm, where sexes can be identified in the caterpillars (Zhang *et al.,* 2010), no strong difference was evident in the global β-diversity structure of the bacterial microbiota. However, marginal differences in the relative abundances of some bacterial taxa was reported, as females were shown to preferentially harbor *Delftia, Aurantimonas* and *Staphylococcus* while males were mostly colonized by *Enterococcus* (Sun *et al.,* 2016). In adult *H. erato,* males and females share similar microbial communities (Hammer *et al.,* 2014), whereas in *Spodoptera littoralis* the sexes harbor divergent bacterial communities, with higher Enterobacteriaceae proportion found in females (Chen *et al.,* 2016). It is noteworthy that even when found, the consequences of sex-dimorphic microbiota in Lepidoptera are not well understood. Chen et al. (2016) showed enrichment of bacteria carrying genes involved in the energetic metabolism in females. Some of these bacterial taxa colonizing females were partly retrieved from the eggs. Those bacteria may be vertically transmitted from the mother to their eggs.

We found that the parasitoid infection was also associated with lower occurrence probability of some taxonomical groups, such as Clostridia, Rhizobiales, Neisseriales and Burkholderiales. This may result from parasitoid infection modifying host’s immune (Tan *et al.,* 2018) or metabolic (Potter and Woods, 2012; Mrinalini *et al.,* 2015) homeostasis that can further influence the intestinal microbial community. Several studies have recently reported an impact of polydnaviruses injected in the caterpillars through the venoms of parasitoid wasps (Cusumano *et al.,* 2018; Tan *et al.,* 2018; Zhu *et al.,* 2018). These symbiotic viruses induce changes in the caterpillar-plant interactions as well as in host immunity. Even though it has never been specifically studied, these viruses might also directly or indirectly impact the microbiota of the caterpillar. Alternatively, individuals not carrying specific symbionts may be more attractive or susceptible to the parasitoid infection. Such processes have been described, for example, in aphids where facultative symbionts interfere with the volatile signals released by the plant to attract parasitoid (Frago *et al.,* 2017). *Wolbachia* sp., on the contrary, were more likely to occur in the gut of parasitized individuals. Previous screening of *M. cinxia* adults have not found presence of *Wolbachia*, whereas the parasitoid, *H. horticola,* is naturally infected by a *Wolbachia* strain wHho, with an infection rate of approximately 50% in the study population the Åland islands (Duplouy *et al.,* 2015). Therefore, our results suggest that *Wolbachia* may be horizontally transferred by the parasitoid. However, due to the high mortality of individuals to the parasitoid infection it may be extremely rare to find Wolbachia infected adults. Furthermore, we do not know whether *Wolbachia* is able to persist in the individuals across the development or if they are viable within the caterpillar gut. As recently reported only 16.3% of the Lepidopteran caterpillars are infected by *Wolbachia* with different impacts of the endosymbiotic bacteria on the reproduction and the sex ratio of their host *e.g.* male killing, feminization, and cytoplasmic incompatibility (Duplouy and Hornett, 2018).

### Effects of host plant’s microbiota and metabolite composition

The microbiome of plant phyllosphere is partially conserved across species with presence of recurrent taxa such as *Methylobacterium, Pseudomonas* and *Sphyngobium* (Delmotte *et al.,* 2009). However, the plant microbiome is also generally considered highly variable and subject to spatial and temporal fluctuation in response to several abiotic factors (Lindow, 1996; Turner *et al.,* 2013). In addition, biotic factors, such as plant genotype, developmental stage or chemical composition are known to affect the microbiome (Delmotte *et al.,* 2009; Berlec, 2012; Bodenhausen *et al.,* 2014; Gargallo-Garriga *et al.,* 2016; González-Arenzana *et al.,* 2017). Consistently with previous results, we showed that the bacterial community of *P. lanceolata* is highly conserved and dominated by a set of core microorganisms, mainly OTUs classified as *Methylobacterium,* that are present in the majority of the samples. These epiphytic Alphaprotebacteria are particularly adapted to the plant phyllosphere and recycle parts of the metabolites secreted by the stomata (methanol and amino acids), and contribute to plant quality, growth and defense (Sy *et al.,* 2005; Madhaiyan *et al.,* 2006; Kutschera, 2007; Madhaiyan *et al.,* 2015).

When considering the whole community structure of the plant microbiota, most of the taxa where correlated with the metabolite profile of the host plant, so that the microbiota tended to decrease with decreasing carbohydrates and amino acid residues. This suggests that these plant metabolites either drive the bacterial communities that successfully colonize the leaves or that the leaf bacterial communities impact plant metabolism. Somewhat surprisingly, the defensive compounds (iridoid glycosides and verbascoside) showed little variation and was not influenced by the plant microbiota.

In general, the microbial communities of host plants and caterpillars were very different. The predominant bacteria in the plants, such as *Methylobacterium* sp., *Hymenobacter* sp., *Modestobacter* sp. and *Aureimonas* sp., were not dominant or even prevalent in the caterpillars. However, a high abundance of OTUs in the host plant did positively affect the same OTUs in the caterpillars in at least few taxonomic groups: in Methylobacteriaceae and some other Alphaproteobacteria, high abundance in the host plant increased their occurrence probabilities in the caterpillars, and in Cytophagaceae and some Methylobacteriaceae, high abundance in the host plant increased the OTU abundance in the caterpillars.

We suggest three potential reasons explaining the observed poor correspondence between caterpillars and host plant microbiota and/or metabolite composition. First, despite the high variability, the bacterial taxa associated with *M. cinxia* gut may be well adapted to their host and consequently little impacted by food intake, including the variation in secondary metabolites, such as iridoid glycosides and verbascoside concentrations. Second, the observed caterpillar gut microbiota variability might reflect high abundance of transient bacteria, which are rapidly acquired and eliminated with high turnover. Third, the microbiota of diapausing caterpillars may shift quickly in the beginning of the diapause, in the absence of plant microbial load or metabolites ingested. Fourth, several species of Lepidoptera harbor horizontally acquired bacterial genes that detoxifies plants secondary metabolites. Such gene acquisitions might have relaxed any selective pressure in favor of the maintenance of bacterial symbionts within the gut leading to high variability of these communities (Hammer *et al.,* 2017; Paniagua Voirol *et al.,* 2018). Our observations are somewhat contrasting with other systems in which nutritionally acquired metabolites of the host plant have been observed to strongly shape the animal gut communities (Koropatkin *et al.,* 2012; Etxeberria *et al.,* 2013; Lu *et al.,* 2014; Xu *et al.,* 2016). Our results also contrast several other studies in insects that have highlighted the importance of host plant in shaping the gut microbiota community (Broderick *et al.,* 2004; Xiang *et al.,* 2006; Pinto-Tomás *et al.,* 2011; Gayatri Priya *et al.,* 2012; Mason and Raffa, 2014; Berman *et al.,* 2018), including a study of actively feeding late instar stage of *M. cinxia* (Ruokolainen *et al.,* 2016). The microbiota of actively feeding individuals are evidently affected by the plant material that they feed on, which can lead to rapid and reversible changes in the microbiota community depending on the organic matter, defensive metabolites. In actively feeding caterpillars the microbiota found in fecal samples has been shown to resemble that of the host plant (Hammer *et al.,* 2017). Our result of microbiota in the midgut not representing similar microbial community to that of the host plant suggest that the bacterial community of the host plant is actively transported through the digestive tract of the caterpillar while they are eating plant material, and that this community is excreted through the feces and is not maintained within the gut of the caterpillar after they stopped eating. On the other hand, we cannot exclude the hypothesis that the excretion of the microbiota have happened during the molting right before the individuals enter into diapause. A recent study on several *Lycaenid* butterfly species showed that starved carnivorous or herbivorous caterpillars did not present any differences in their intestinal communities in comparison to each other (Whitaker *et al.,* 2016).

### The over-winter survival probability of caterpillars families is spatially structured but does not correlate with the microbiota or metabolite composition of the host plant

We did not find any influence of microbiota composition on overwinter survival, but this may have been because our data on this was indirect. As the microbiota of the caterpillars from the same nest (i.e. family) did not resemble much each other, the microbiota of the sampled individuals was not likely to be representative of the microbiota of the individuals for which we scored survival. Previous studies on Lepidoptera have documented contradictory results on the impact of gut microbiota on survival. Experimental perturbation of *Manduca sexta* microbiota by antibiotic treatments had no effect on survival and development (Hammer *et al.,* 2017), whereas the removal of *Enterococcus munditii* symbionts colonizing *Galleria mellonella* decreased individual survival during the adult stage (Johnston and Rolff, 2015). The observed over-winter survival of the *M. cinxia* families in the wild exhibited some spatial structure, suggesting that the mortality is strongly influenced by some spatially autocorrelated environmental factor such as summer drought (Saastamoinen *et al.,* 2013; Tack *et al.,* 2015, Kahilainen et al. 2018) or host plant density, not accounted for in our study.

### Conclusion

The caterpillars of the Glanville fritillary butterfly present a highly variable gut microbiota even among caterpillars from the same family living on the same host plant individual. Variation in gut microbiota is predominantly related to Enterobacteriaceae, which show marked variation in their diversity among the individuals. Additionally, the occurrence probabilities of some OTUs were impacted by the presence of the parasitoid and by the sex of the caterpillar. The highly variable herbivore microbial communities differed markedly from those of the more conserved host plant microbiota communities. In particular, while the plant leaf metabolites influenced the plant microbiota, these effects did not penetrate to microbiota of the caterpillars feeding on those leaves. Future prospects on other developmental stages (pupae, adults, eggs) should be conducted to broaden our understanding of the variation and potential role of the Glanville fritillary microbiota. Finally, we remark that joint species distribution modelling provides new opportunities to assess simultaneously the influences of both abiotic and biotic drivers on community variation. However, further development is required for making joint species distribution models fully compatible with sequencing data, and thus we hope future developments to implement multinomial observation models and the ability to capture discrete mixtures.

## Experimental Procedures

### The study system

The Glanville fritillary, *Melitaea cinxia,* butterfly occurs across the Eurasian continent, and in northern Europe has a univoltine life-cycle (Ehrlich and Hanski, 2004). In Finland, the butterfly occurs only in the SW archipelago, the Åland islands, where it persists as a classic metapopulation within a network of ~4.000 discrete habitat patches consisting of meadows and pastures (Ojanen *et al.,* 2013). The habitat patches have been annually surveyed since 1993 for the presence of caterpillar family nests (Hanski, 1994; van Nouhuys and Hanski, 2005; Ojanen *et al.,* 2013). Females lay clutches of eggs on two caterpillars host plant species, *Veronica spicata* and *Plantago lanceolata* (Kuussaari *et al.,* 2000). The gregarious caterpillars develop within the host plant, and in the fall, they build a thick and conspicuous winter nest, terminate feeding and molt into diapausing morphotype (Wahlberg, 2000). The diapause is broken in spring when the caterpillars continue their development until pupation. Approximately, 30% of the caterpillar families die during the winter (Tack *et al.,* 2015). In addition, a conserved proportion of approximately 30% of the individuals get infected by a specialist parasitoid *Hyposoter horticola* (Ehrlich and Hanski, 2004; Nouhuys and Ehrnsten, 2004). The parasitism occurs during the egg stage, after which the parasitoid develops within the host until it hatches from the 7^th^ instar caterpillars early in the spring and kills the host. Several reasons make this system suitable for the present study: (i) the caterpillars and their host plant can be easily found from the field due to the gregarious life-history of the caterpillars and the conspicuous silk nest they spin for over-wintering; (ii) the over-wintering caterpillars are synchronized in their development prior diapause and have an empty gut at this developmental stage (Ojanen *et al.,* 2013), which reduces confounding factors in the analyses; (iii) several individuals, from mainly full-sib families (Fountain *et al.,* 2018), can be sampled from the same over-wintering nest on one host plant individual, which allows us to assess individual variation both within and among families; (iv) the local populations are well-described due to the long-term ecological monitoring; and (v) the host sex can be identified at the caterpillar stage using molecular markers (Rastas *et al.,* 2013).

### Sample collections

Caterpillar and plant samples were collected from natural populations of the *M. cinxia* in the region of Sund in the Åland islands within three-day period in September 2015. This region was selected due to generally high occupancy of the butterfly in three connected networks ensuring sample availability (Supplementary material), and the possibility to control for some potentially confounding factors due to dominance of only one host plant species (*P. lanceolata)* and one specialist parasitoid species (*H. horticola)* (Nair *et al.,* 2016; Hanski *et al.,* 2017). The survey followed the general framework of the long-term survey of the *M. cinxia* butterfly (described in Ojanen *et al.,* 2013: a total of 189 dry meadows i.e. potential habitats) were surveyed for the presence of winter nests. Once located, the GPS coordinates were registered using the Earthcape biodiversity platform (http://www.earthcape.com). From each nest, three 5^th^ instar caterpillars and one leaf from the host plant on which the caterpillars resided were collected with disinfected forceps and stored individually in sterile 1.5 ml and 15 ml tubes, respectively. A total of 191 caterpillars from 66 nests and 63 host plant samples were collected from the 15 patches that were occupied by the butterfly in 2015. In few cases, the entire host plant had already been consumed, and hence no plant sample was collected. The caterpillars were dissected in order to detect the presence of the potential parasitoid and to separate midgut from rest of the carcass (for more details about sample conservation and preparation see Supplementary Material). Insect digestive tract is separated in three sections (foregut, midgut, hindgut) with often observed heterogeneity in their physiology but also in the composition of the microbial communities (Engel and Moran, 2013). Results on Lepidoptera have, however, been somewhat contradictory, with differences in the microbiome across the different gut sections being evident in *Spodoptera litteralis* (Tang et al., 2012) but not in *Bombix mori* (Chen et al., 2018). To avoid merging communities that potentially differ, we focused specifically on the microbiota localized within the midgut of the caterpillars. This section is the largest section, most important for food digestion, and its microbiota often shows interaction with host plant secondary metabolites (Terra and Ferreira, 2012; Pentzold *et al.,* 2014). The over-winter survival of caterpillars nests in the field (i.e. from which the three individuals were sampled from) was assessed in the spring 2016, by checking the presence of active post-diapause caterpillars (Ojanen *et al.,* 2013).

### High throughput rrs amplicon sequencing

DNA was extracted from midgut samples with Qiagen DNeasy Blood and Tissue kit (Qiagen, Germany) using an optimized protocol for extraction of bacterial DNA from low matrix (Minard *et al.,* 2015). For plant samples, a piece of 0.5 cm^2^ was separated from the center of the leaf, crushed in liquid nitrogen using a sterile pestle, and DNA was extracted following the protocol described for midgut samples. To avoid bias due to the possible confounding effect of extraction set, the samples were randomized before extraction. In addition, three independent extractions were carried out without any matrix and processed with the rest of the samples to identify potential bacterial DNA contamination that could affect results obtained from low biomass samples (Salter *et al.,* 2014).

The 280bp hypervariable V5-V6 region of the *rrs* gene was amplified in duplicates and sequenced with Miseq v.3. sequencing platform (Illumina, U.S.A.). Details on the protocol are available in the Supplementary material. Analysis of sequences was performed using mothur v.1.37.6 following the Miseq Standard Operating Procedure described by the developers (http://www.mothur.org/wiki/MiSeq_SOP) (Schloss *et al.,* 2009). A total of 16,710,206 sequences were obtained after alinement of forward and reverse reads. Aligned sequences were selected within a size range of 250-350 bp with less than 8 homopolymers and any ambiguous position. All sequences which did not align to the *rrs* Silva v.123 database were filtered out. *De novo* chimera detection was performed using UCHIME implemented in mothur (Edgar *et al.,* 2011). Clustering was performed using a maximum of 3% distance within each Operational Taxonomic Unit (OTU) according to the average neighbor method. After quality trimming and clustering, every contaminant sequence was trimmed out from the Sample x OTU shared table as previously described (Minard *et al.,* 2015). The samples were first rarefied at 3000 reads in order to control for sequencing depth biases. Same OTUs were considered as contaminant if they were present in the negative controls and if their proportion in a given sample was not at least 10 times higher than their proportion in the negative controls. After trimming and quality control, the samples were rarefied at 1500 reads per sample for further analysis. 20 caterpillar samples and 2 plant samples, which did not contain the minimum amount of sequences, were discarded for the rest of the analysis. Miseq sequences have been deposited on the European Nucleotide Archive (http://www.ebi.ac.uk/ena) under the accession project number PRJEB26629.

### Metabolomic analysis of the leaf samples of host plant Plantago lanceolata

After subtraction of the extremities, the remaining parts of each leaf sample were crushed with a sterile pestle in liquid nitrogen and the frozen powder was freeze dried for 48h. The extraction was processed using previously described protocol and ^1^H-NMR spectra of the metabolites were recorded (Supplementary Material; Kim *et al.,* 2010). NMR spectra were processed with MNOVA software v.10.0.2 (Mestrelab research S.L., Spain). Model compounds of Aucubin (Sigma-Aldrich, Germany), Catalpol (Sigma-Aldrich, Germany) and Verbascoside (Extrasynthese, France) were used for signal assignments of *P. lanceolata* defensive metabolites. Other primary or secondary metabolite shifts, and J-coupling constants obtained from plant material using similar solvents were used as reference (Kim *et al.,* 2010; Lubbe *et al.,* 2011; Yang *et al.,* 2012; Agudelo-Romero *et al.,* 2014; Gallo *et al.,* 2014). For multivariate analysis, the signal was binned to 0.04 ppm and integrated. The TSP and Methanol signals were removed and the relative intensity of the chemical signals were normalized according to the dry mass of the samples and the TSP intensity.

### Sex determination

As caterpillar’s sex cannot be determined based on morphology, we employed a panel of 24 SNP markers linked to the Z chromosome to differentiate the sexes (Supplementary material, Supplementary Table S3, S4). The sensitivity and specificity of this method was estimated to be 0.81 and 0.89, respectively, based on a group of 150 adult individuals with known gender (75 males and 75 females). A total of 15 individuals could not be annotated based on the SNP panel.

### Statistical analyses

We analyzed the data with Hierarchical Modelling of Species Communities (HMSC; Ovaskainen *et al.,* 2017), which approach belongs to the class of joint species distribution modeling (JSDM; Warton *et al.,* 2015). HMSC provides simultaneously species- and community-level inference on how species occurrences and/or abundances relate to environmental covariates, and how these relationships are structured with respect to species traits and phylogenetic relationships. HMSC additionally assesses the structure of co-occurrence patterns among the species that can’t be attributed to responses of the species to the measured covariates, either in spatially hierarchical or in spatially explicit context, depending on the nature of the study design (Ovaskainen *et al.,* 2017).

We performed two separate analyses, called hereafter caterpillar and plant models, which differed in whether the OTU data were derived from caterpillar or plant material. In both models, the response variable was the vector of rarified sequence counts of the microbial OTUs. We employed a hurdle approach, in which we first used a probit model for OTU presence-absence, and then a log-normal model for OTU abundances conditional on presence. We restricted the analyses to OTUs that were present in at least five samples (562 and 610 OTUs for caterpillars and plants, respectively). We further excluded samples for which plant OTUs or metabolites were missing. The analyzed dataset consisted of 142 caterpillars collected from 55 host plants (Fig. 2).

In the caterpillar model, our aim was to examine how the OTU composition depended on the properties of the focal caterpillar, and on the OTU and metabolite compositions of its host plant. We included as fixed effects (1) the sex of the individual (0 for female and 1 for male), (2) the infection status of the individual (0 for non-infected and 1 for infected by the parasitoid wasp), (3) the abundance of the focal OTU in the host plant where the individual was residing, (4) the plant OTU community composition, and (5) the plant metabolite composition. We measured plant OTU abundance as log-transformed sequence count and described plant OTU community composition and plant metabolite composition by the first three principal components that explained respectively 22% and 92% of their total variations (Supplementary Fig. S3, S7). To examine whether the responses of the species to the explanatory variables showed a phylogenetic signal, we included in the analysis a phylogenetic correlation matrix among the OTUs, obtained with FastTree method assuming the General Time Reversible (GTR) evolution model (see Supplementary Fig. S8) (Price *et al.,* 2010). To examine residual co-occurrence patterns among the OTUs that cannot be attributed to the fixed effects, we further included in the model the level of the caterpillar nest (i.e. host plant level) as a spatial random effect, and the level of the individual caterpillars as a non-structured random effect. In the plant model, we included as the sole fixed effect the plant metabolite composition, and as the sole random effect the level of the level of the plant as a spatial random effect.

We fitted both the caterpillar and the plant models using the HMSC-Matlab implementation of Ovaskainen et al. (2017) with default prior distributions. To examine how much of the variation in OTU occurrences can be attributed to the fixed effects and to associations among the OTUs, we evaluated the predictive power of the model in three different ways. All of these accounted for the fixed effects, but differed on how the random effects were accounted for. Prediction P1 is aimed at measuring the predictive power based solely on fixed effects, and thus we integrated the random effects over their prior distributions rather than using sampling unit -specific fitted values. Prediction P2 is aimed at measuring the predictive power that can be gained by accounting for species-to-species associations. To generate P2, we split the species randomly to two groups, and made the predictions for each species group conditionally on the known occurrences of species belonging to the other group (see Supporting Information for details). Prediction P3 is aimed at measuring the full explanatory power of the model, and thus here the random effects were included based on their fitted values. Therefore, the performance of P1 measures the importance of fixed effects, and the difference between P2 and P1 (respectively, between P3 and P1) gives a minimum (respectively, maximum) estimate for the importance of species-to-species associations. This is because the difference between P3 and P2 may either be a true effect of species-to-species associations that is not captured by our approach of dividing the species into two groups, or then it may be due to overfitting of the random effects. We measured predictive powers by Tjur’s R^2^ (Tjur, 2009) for the probit models and standard R^2^ for the log-normal models. Given that HMSC framework has not previously been used in microorganism studies and may thus not be familiar to microbial scientific community, we ran a series of complementary analyses with more traditional methods to support our HMSC-based results. The details are given in Supplementary material.

Finally, we analyzed whether the overwintering survival of caterpillar nests (siblings of the caterpillars assessed above) was dependent on metabolite and OTU composition of the host plant they were residing on. We performed this analysis with a logistic regression model estimated with STAN (Carpenter *et al.,* 2017), in which model we accounted for the spatial locations of the nests using a Gaussian process approach (see Supplementary material) (Rasmussen and Williams, 2006).

## Acknowledgements

Funding for this project was provided by grants of the European Research Council (Independent Starting grant no. 637412 ‘META-STRESS’ to MS) and the Academy of Finland (Decision numbers 273098 and 265641 to MS and 1273253, 250444 and 284601 to OO) and the Research Council of Norway (CoE grant 223257), as well as by the Biocenter Finland related to the NMR core facility at the Institute on Biotechnology. We acknowledge Juha-Matti Pitkänen for help with DNA extraction, Sami Ojanen for the coordination of sampling, and field assistants for sample collection. We would also like to thank Aapo Kahilainen for his help with genotyping analysis.

## Conflict of Interest

The authors declare no conflict of interest.

