## Supplementary material for "Variation in *Melitaea cinxia* gut microbiota is phylogenetically highly structured but only mildly driven by host plant microbiota, sex or parasitism": Supp_legend

**SUPPLEMENTARY FIGURE LEGENDS:**

**Supplement Figure S1. Results of Dirichlet-multinomial analysis.** Panels A and B depict the Laplace goodness-of-fit measures for fitted mixture models with different number of mixture components (lower values corresponds to better fit). Panel C visualize the 2 mixture components of the best model for microbial OTUs in caterpillars.

**Supplement Figure S2. Results of permutation tests.** Each panel depict the Spearman rank correlation coefficient between the assigned mixture component of the best 2-component Dirichlet-multinomial model for caterpillar OTUs data and available predictors. Red line corresponds to the real value and the black curve depict the density of permutation-based values. Dashed blue lines depict the 2.5% and 97.5% quantiles of the permutation-based density.

**Supplement Figure S3. Principal Component Analysis (PCA) of the metabolites associated with the host plant.** The PCA plot represents the ordination of the plant metabolites on the three first Principal Components (A) PC1 and PC2, (B) PC1 and PC3, (C) PC2 and PC3. The signal corresponding to the chemical shift of carbohydrates and amino acid residues are colored in red while other signals are colored in blue.

**Supplement Figure S4. Partitioning of the explained variance of bacterial OTUs among the fixed and random effects in plant models.** The colored bars show, for each OTU, the proportions of variance attributed to each of explanatory variables. The average variance proportions over the OTUs are shown in the legend box. The ordering of OTU is following ordering of Fig. 1 except for the OTUs that were recorded only in larvae samples (for details, see Supplementary Table S2). See *Statistical Methods* for a full description of the included fixed and random effects.

**Supplement Figure S5. The influence of metabolic covariates on plant microbiota.** Regression coefficients that were estimated to be positive (respectively, negative) with 95% credibility level are shown by red (respectively, blue). The ordering of the OTUs is identical to that of Fig. S4.

**Supplement Figure S6. Residual associations among plant microbiota.** The panels illustrate the random effects for the presence-absence (A) and abundance (B) parts of the plant model. OTU-pairs for which the residual correlation was estimated to be positive (respectively, negative) with 95% credibility level are shown by red (respectively, blue) color. The ordering of the OTUs is identical to that of Fig. S4.

**Supplement Figure S7. Principal Component Analysis (PCA) of the bacterial microbiota associated with the host plant.** The PCA plot represents the ordination of the bacterial Operational Taxonomic Units (OTUs) on the three first Principal Components (A) PC1 and PC2, (B) PC1 and PC3, (C) PC2 and PC3. The color scale represents the OTU classification at the Phylum level.

**Supplement Figure S8. OTUs phylogenic relationship matrices.** Phylogenetic relationship among OTUs are represented for larvae (A) and plant (B) microbial The relationships between the OTUs, used for analysis of bacterial community, were obtained with FastTree method assuming the GTR evolution model. Color of each cell encodes the relationship between the OTUs, located at those row and column with the gradation of red indicating the level of relatedness. The order of the OTUs is selected according to the available approximate taxonomic classification and further aligned according to phylogenic similarity, with the colors and relative ordering following the Fig. 2 in main text. A detailed similarly ordered lists of individual OTU with their full taxonomic classification are provided in Supplementary Table S2.

**SUPPLEMENTARY TABLE LEGENDS:**

**Supplementary Table S1. Predictive powers of the larval and plant models.** Predictive power is measured by Tjur R^2^ for the presence-absence models and by the standard R^2^ for the abundance models. The values show the mean ± standard deviation over the OTUs. As detailed in the *Statistical Methods,* Prediction P1 measures the predictive power solely due to the fixed effects part of the models, whereas P2 and P3 also account for species-to-species associations, with P2 being based on cross-validation across species and P3 in on predicting the same data that were used to fit the model.

**Supplementary Table S2. Taxonomic classification of the bacterial Operational Taxonomic Units.**

**Supplementary Table S3. Proportion of Heterozigous loci observed in males and females within the validation sample panel with known gender.**

**Supplementary Table S4. Genotyping for sex determination of *M. cinxia*.**
