## Supplementary material for "Variation in *Melitaea cinxia* gut microbiota is phylogenetically highly structured but only mildly driven by host plant microbiota, sex or parasitism": supp_Methods

**Supplementary Material and Methods**

**Conservation of samples and preparation of larvae**

During the field collection period, larval samples and plant samples were stored in a liquid nitrogen tank and a -20°C freezer, respectively. Samples were transported in dry ice to the laboratory and stored in a -80°C freezer until processed. Before processing, the larval surface was disinfected. Each larva was washed three time with 1X Phosphate Buffer Saline (Ambion, U.S.A), then immerged in 70% ethanol for 5 minutes and rinsed 5 times with 200µl of 1X Phosphate Buffer Saline. The larvae were dissected in 1X Phosphate Buffer Saline within a sterile environment. The gut was separated from the carcasses and the midgut was isolated. For each individual, the haemocoel was dissected to identify a potential infection by the specialist parasitoid *H. horticola.* Once identified, the parasitoid was separated from the carcass.

**V5-V6 amplification and sequencing of larvae midguts and the host plant leaves**

For each sample with 3µl of the DNA extract and modified primers 784F (5’-AGGATTAGATACCCTGGTA-3’) and 1061R (5′-CRRCACGAGCTGACGAC-3′) containing Illumina sequencing primers (Illumina, U.S.A.) (Andersson *et al.,* 2010). Amplifications of the fragments were performed in duplicates with 1 X of Q5 reaction buffer (New England Biolabs, U.S.A.), 1X of Q5 High GC enhancer (New England Biolabs, U.S.A.), 0.5 unit of Q5 High-Fidelity DNA Polymerase (New England Biolabs, U.S.A.), 0.04 mM of dNTP (New England Biolabs, U.S.A.), 0.2 µM of each primer, 8 µg.µl^-1^ of Bovine Serum Albumine (New England Biolabs, U.S.A.), 1.2 µg.µl^-1^ of T4gene32 (New England Biolabs, U.S.A.) and 3 µl of DNA (19.6 ± 1.94 ng). The amplifications were conducted as previously described (Minard *et al.,* 2015). The duplicate products were pooled per individual samples. The pools were purified and indexed before sequencing by the Institute for Molecular Medicine Finland (FIMM, Finland) using the Miseq v.3. sequencing platform (Illumina, U.S.A) to a read length of 300bp paired-end in order to fully cover the V5-V6 region with both forward and reverse pairs. To increase the genetic diversity and improve the cluster separation, 5% PhiX phage DNA was added to the libraries prior sequencing.

**Metabolites extractions from the host plant leaves and acquisition of the ^1^H-NMR spectra**

The extraction was processed using the previously described protocol (Kim *et al.*, 2010). Metabolites were extracted from 3.5 mg of dried powder of each sample using a solvent containing 750 µl of CD_3_OD (VWR Chemicals, Belgium) and 750 µl of KH_2_PO_4_ (Sigma-Aldrich, Germany) buffer in D_2_O (pH6) with 0.05% (wt/wt) of TSP (sodium trimethylsilylpropionic acid) (Sigma-Aldrich, U.S.A.). The mixture was sonicated for 20min and then centrifuged at 17,000g for 10min. The supernatant was collected and 800 µl was transferred into a 5mm Ø NMR tube (Wilmad, U.S.A.). Proton Nuclear Magnetic Resonance (^1^H-NMR) spectra were acquired at 298K on a Bruker 850 MHz Avance III HD NMR spectrometer equipped with a TCI Cryoprobe (Bruker, U.S.A.). ^1^H NMR spectra were recorded using 1D presaturation pulse sequence (zgpr). For each ^1^H spectrum, 256 transients were collected into 32K time domain points using a 60° flip angle, spectral width of 10.2 kHz, relaxation delay of 5.0s, an acquisition time of 1.6s, and a mixing time of 5ms. Fourier transformation of the free-induction decay was applied with zero filling to give 65K frequency domain data points. Prior to Fourrier transformation, 0.3 line broadening was applied. ^1^H chemical shifts were referenced to an internal standard TSP at δ 0.0 ppm.

**Sex determination**

DNA was extracted from larvae carcasses using nucleospin 96 tissue spin kit (Macherey-Nagel, Germany) and quantified with Qubit dsDNA HS Assay Kit (Life technologies, USA). SNP genotyping was performed with Sequenom iPLEX Gold (Sequenom inc., USA) chemistry at the Institute for Molecular Medicine Finland (FIMM, Finland) using 120ng of DNA. A total of 11 samples were genotyped in duplicate (at least one duplicate per plate) as well as 4 water controls. In Lepidoptera, individuals with a Z0, ZW or ZWW chromosome karyotype are females and homogametic individuals with a ZZ karyotype are males (Traut and Marec, 1997). The individuals were considered as males if at least two SNPs presented a heterozygous pattern reflecting the presence of two Z chromosomes copies. A total of 23 loci only associated with the Z chromosome were used in this experiment. In addition, one pseudoautosomal loci (present in both W and Z chromosomes and for which females can be heterozygous) was also used to improve the accuracy of the assignment. The targeted loci as well as their heterozygosity proportion in males and females (obtained from the panel of 150 adult individuals with known gender) have been summarized in the Tables S3 and S4.

**Statistical analyses**

*Larvae bacterial model*

We analyzed the larvae OTU data with Hierarchical Modelling of Species Communities approach (HMSC; Ovaskainen *et al.*, 2017), and exploited the hurdle approach to account for potential zero-inflation of the rarified sequence counts: first we fitted the probit model to the presence-absence of OTU in larvae, and then we fitted the lognormal model based only on those observations, where the OTUs were present. We denote by $n_{y}=142$ the number of observed larvae, by $n_{p}=55$ – number of different host plants that the larvae were residing, by $n_{s}=562$ the number of studied OTUs, by $n_{c}=10$ the number of included univariate covariates to the model, by $n_{f}^{p}$ – the estimated number of latent factors at the plant random effects level, and by $n_{f}^{l}$ – the estimated number of latent factors at the larvae random effects level.

In both probit and lognormal models the linear component of the model was defined as

$$L=XB+{\Pi H}_{p}\Lambda_{p}+H_{l}\Lambda_{l},$$

where $L$ – matrix of latent liabilities of size $n_{y}\times n_{s}$, $X=\left[ X_{I},X_{p},X_{s},X_{f},X_{o_{1}},X_{o_{2}},X_{o_{3}},X_{m_{1}},X_{m_{2}},X_{m_{3}} \right]$ – combined matrix of included covariates with size $n_{y}\times n_{c}$, $B$ – matrix of OTUs’ responses to covariates, $H_{p}$ and $\Lambda_{p}$ – matrices of latent factors and latent loadings at the plant level of sizes $n_{p}\times n_{f}^{p}$ and $n_{f}^{p}\times n_{s}$ correspondingly, $\Pi$ – a matrix defining the projection of larvae to its host plant of size $n_{y}\times n_{f}^{p}$, $H_{l}$ and $\Lambda_{l}$ – matrices of latent factors and latent loadings at the larvae level of sizes $n_{y}\times n_{f}^{l}$ and $n_{f}^{l}\times n_{s}$ correspondingly. The covariate matrix consisted of

1. $X_{I}$ – the intercept, vector of ones.
2. $X_{p}$ – the parasitoid infection status of larvae individuals, equal to 1 iff the larvae was infected.
3. $X_{s}$ – the gender of the larvae, equal to 1 if male, 0 if female. For those larvae, where the sex determination failed we substituted the value in this column with 0.5.
4. $X_{f}$ – an OTU-specific column, containing the zero-corrected log-abundance of the focal species $j$ in the host plant.
5. $X_{o_{1}},X_{o_{2}},X_{o_{3}}$ – the first three principal components of the bacterial community in the host plant.
6. $X_{m_{1}},X_{m_{2}},X_{m_{3}}$ – the first three principal components of metabolic profile in the host plant.

Additionally, the phylogenic relationship matrix was included to account for relatedness between the OTUs in the community through imposing the similarity on single OTUs responses to covariates (columns of matrix $B)$. Latent factors on the larvae level were assumed non structured and on the plant level assumed to be spatial. In probit model the data distribution was specified as $Y_{ij}\sim Bernoulli\left( probit^{-1}\left( L_{ij} \right) \right)$, where $Y_{ij}$ is equal to 1 iff OTU $j$ was found to be present in $i$-th larvae. In lognormal model the data distribution was specified as $\log\left( Y_{ij} \right)\sim N\left( L_{ij},\sigma_{j} \right)$, where $Y_{ij}$ is the abundance of OTU $j$ in the $i$-th larvae, and $\sigma_{j}$ is the OTU-specific dispersion parameter. All model parameters were assigned default HMSC priors.

*Plant bacterial model*

In plant analysis we employed approach, very similar to our analysis of larvae. We denote by $n_{y}=55$ the number of observed larvae, by $n_{s}=610$ the number of studied OTUs, by $n_{c}=4$ the number of included univariate covariates to the model, by $n_{f}$ – the estimated number of latent factors at the plant random effects level The linear model component was specified as

$$L=XB+H_{p}\Lambda_{p},$$

where $L$ – matrix of latent liabilities of size $n_{y}\times n_{s}$, $X=\left[ X_{I},X_{m_{1}},X_{m_{2}},X_{m_{3}} \right]$ – combined matrix of included covariates with size $n_{y}\times n_{c}$, $B$ – matrix of OTUs’ responses to covariates, $H_{p}$ and $\Lambda_{p}$ – matrices of latent factors and latent loadings at the plant level of sizes $n_{y}\times n_{f}$ and $n_{f}\times n_{s}$ correspondingly. The covariate matrix consisted of

1. $X_{I}$ – the intercept, vector of ones.
2. $X_{m_{1}},X_{m_{2}},X_{m_{3}}$ – the first three principal components of metabolic profile in the plant.

All model parameters were assigned default HMSC priors.

*Predictions and R^2^*

We generated the model predictions in three ways. In all three cases we used the model fitted to the whole dataset, and then varied the underlying assumptions on what parts of information on the fitted model is available for making predictions. First, we simply evaluated how closely the model fits to the training data, thus exploting both the information on the fixed and random effects included to the model. Second, we evaluated how much of predictive power is attributed to the fixed effects of the model, and thus could be considered as an optimistic expectation for predicting new data that was not used in training. Third, to examine how much the estimated co-occurrences among the species influenced the predictive performance of the model, we predicted the occurrences of the OTUs conditional on the data on some of the other OTUs (Ovaskainen *et al.*, 2017). To do so, we split the OTUs in the prediction phase randomly into two equally large groups. We then made the predictions for group 1 conditional on the observed data for group 2, and the predictions for group 2 conditional on the observed data for group 1. Thus we utilized the information on the fixed effects and on the estimated association structure from the fitted model, but assumed no knowledge on the estimated random factors’ distributions.

*Residual checks in lognormal models*

To justify the applicability of lognormal distribution for modelling the abundance data conditional on presence we’ve ran the checks for the residual normality in lognormal models. These checks indicated that out of 562 OTUs in caterpillar model only for 10 (1.8%) the normality hypothesis could be rejected at p=0.05 significance level in Kolmogorov-Smirnov test. In plant model, out of 610 OTUs modeled 29 (4.8%) indicated lack of normality in the residuals with similar test.

*Complementary statistical analysis*

Given that HMSC framework has not been used in microorganism studies so far and is probably not familiar to applied microbial ecologists, we ran a series of complementary analyses with traditional methods to support our HMSC-based results. Specifically, both for caterpillar and plant OTU communitites we exploited the Diriclet-mixture approach, proposed by (Holmes *et al.*, 2012) to test how many of distinct clusters does the data separate to.

As the Diriclet-mixture approach approach indicated that the observed variation in OTUs in caterpillars most plausible under 2 mixture components (Fig. S1A), we tested whether the probability of a caterpillar to be assigned to one of these components could be related to any of the measured caterpillar or host plant characteristics. We used permutation testing to assess, whether the correlations between the assigned mixture component and each of the predictors is plausible under the neutral assumption, or whether such hypothesis is rejected. We made 10^5^ permutations of the data and checked whether the Spearman rank correlation coefficient for the observed data was within [2.5%,97.5%] interquantile range of the correlations for permuted data. Out of 10 predictors in caterpillar model, only for the parasitioid infection the neutral hypothesis was rejected (Fig. S2).

*Survival*

The model structure was $p_{i}=logit^{-1}\left( \beta_{0}+\sum_{k=1}^{n} \beta_{k}^{1}x_{ik}^{1}+\sum_{k=1}^{m} \beta_{k}^{2}x_{ik}^{2}+\eta_{i} \right)$, where $p_{i}$ is the survival probability of a family residing at the plant $i$; $x_{ik}^{1}$ is the value of $k$-th principal component for plant $i$ of metabolic profile principal components decomposition; $x_{ik}^{2}$ is the value of $k$-th principal component for plant $i$ of plant bacterial OTU principal component decomposition; $\eta_{i}$ is the value of spatial effect at plant $i$. Being restricted by relatively small data of 55 binary observations (42 positive outcomes and 13 negative), we limited the choice of explanatory covariates to the first $n=2$ principal components of the metabolic profile and the first $m=2$ principal components of the log-transformed abundance of bacterial OTU community. The spatial effect had a Gaussian process structure $\eta\sim GP\left( 0,k\left( s_{1},s_{2} \right) \right)$ with zero mean and exponential covariance function $k\left( s_{1},s_{2} \right)=\sigma^{2}\exp\left( -\frac{dist(s_{1},s_{2})}{l} \right)$, parametrized by variance $\sigma^{2}$ and range $l$ (Rasmussen and Williams 2006). Additionally, we separately tested the model without the spatial effect. All model parameters were assigned weakly informative priors, that could be considered essentially flat in the plausible region of parametric space.

The model described above was coded in Stan probabilistic programming language using marginal form of the latent variable and non-central Gaussian parametrization (Betancourt et al. 2017):

```
data {
   int<lower=1> Ds;
   int<lower=1> N;
   int<lower=1> M;
   real<lower=0> betaSigma2;
   matrix[N,Ds] s;
   matrix[N,M] X;
   int<lower=0,upper=1> y[N];
}
transformed data {
   matrix[N,N] dist;
   matrix[N,N] SigmaFix;
   for(i in 1:N)
      for(j in 1:N)
         dist[i,j] = pow(dot_self(s[i,]-s[j,]), 0.5);

   SigmaFix = betaSigma2*tcrossprod(X);
}
parameters {
   vector[N] f0;
   real<lower=0> l;
   real<lower=0> s2;
}

model {
   matrix[N,N] K;
   matrix[N,N] L;
   vector[N] f;
   K = s2*exp(-inv(l)*dist);

   f = cholesky_decompose(K+SigmaFix)*f0;
   y ~ bernoulli_logit(f);

   f0 ~ normal(0,1);
   l ~ normal(0,3);
   s2 ~ gamma(1, 0.5);
}
```

This model was fitted using the NUTS sampler (Hoffman and Gelman 2014) with default adaptation parameters and random initiation. We ran 4 chains and 2000 iterations in each chain, out of which first 1000 were used for adaptation and the later 1000 were included to the resulted posterior. The model fit diagnostics indicated proper within and between chain mixing.

None of the included fixed effects were estimated to be sighnificantly below or above zero.

| Posterior distribution quantiles of fixed effects in the survivial analysis | | | | | |
| --- | --- | --- | --- | --- | --- |
|  | Intercept ($\beta_{0}$) | Met_PC1 ($\beta_{1}^{1}$) | Met_PC2 ($\beta_{2}^{1}$) | OTU_PC1 ($\beta_{1}^{2}$) | OTU_PC2 ($\beta_{2}^{2}$) |
| 5% | -0.5545 | -0.3778 | -0.8320 | -1.1857 | -1.0918 |
| 95% | 2.9491 | 1.0990 | 0.5621 | 0.3202 | 0.4709 |
