## Supplementary material for "Variation in *Melitaea cinxia* gut microbiota is phylogenetically highly structured but only mildly driven by host plant microbiota, sex or parasitism": Supp_figures

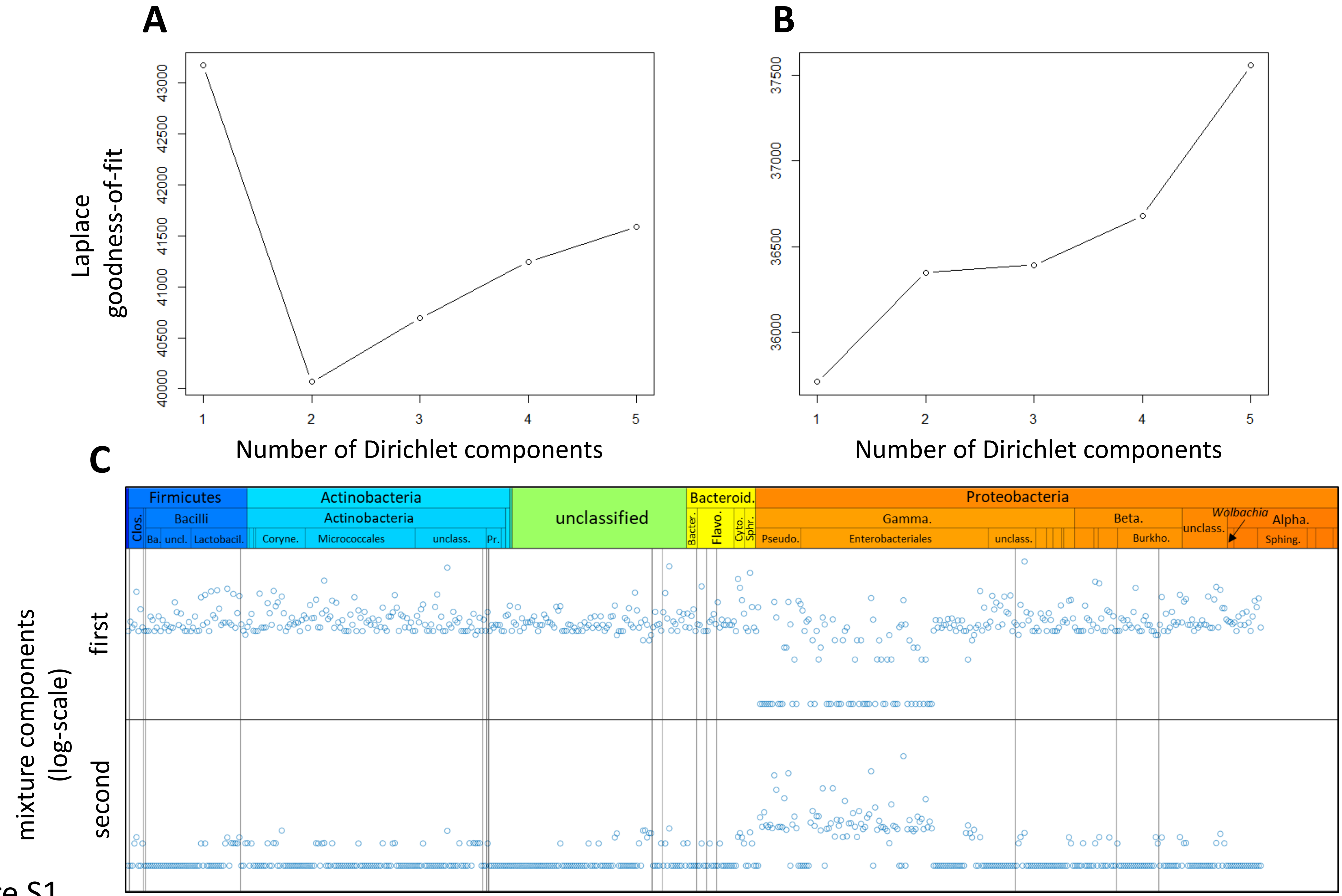
A
B
C
Laplace goodness-of-fit
Number of Dirichlet components
Number of Dirichlet components
mixture components (log-scale)
first
second
Firmicutes
Actinobacteria
unclassified
Bacteroid.
Proteobacteria
Clos.
Bacilli
Ba.
uncl.
Lactobacil.
Actinobacteria
Coryne.
Micrococcales
unclass.
Pr.
Bacter.
Flavo.
Cyto.
Sphr.
Gamma.
Pseudo.
Enterobacteriales
unclass.
Beta.
Burkho.
unclass.
Wolbachia
Alpha.
Sphing.

Figure S1

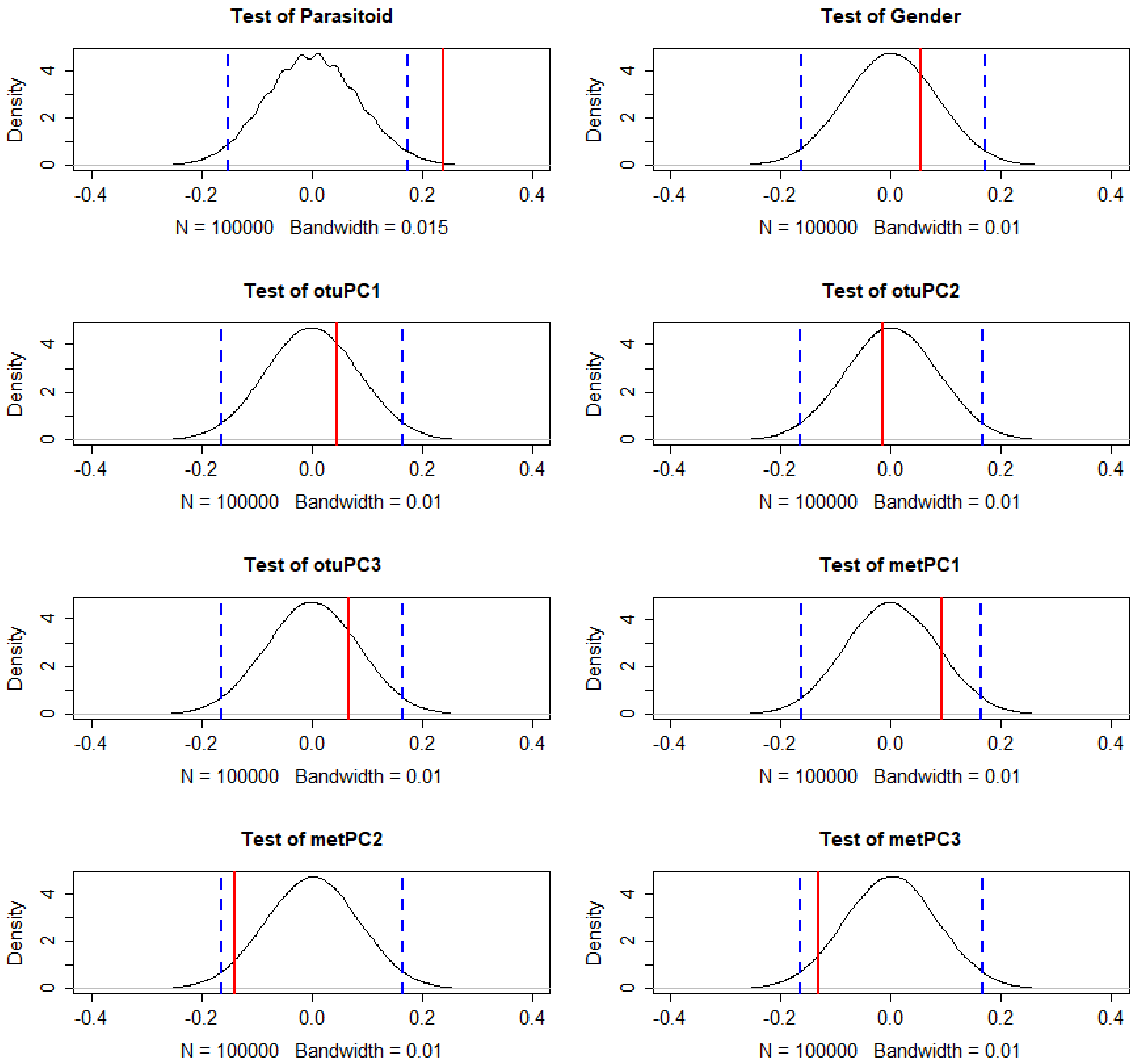
Test of Parasitoid
Density
N = 100000 Bandwidth = 0.015
Test of Gender
N = 100000 Bandwidth = 0.01
Test of otuPC1
N = 100000 Bandwidth = 0.01
Test of otuPC2
N = 100000 Bandwidth = 0.01
Test of otuPC3
N = 100000 Bandwidth = 0.01
Test of metPC1
N = 100000 Bandwidth = 0.01
Test of metPC2
N = 100000 Bandwidth = 0.01
Test of metPC3
N = 100000 Bandwidth = 0.01
-0.4
-0.2
0.0
0.2
0.4
0
2
4

Figure S2

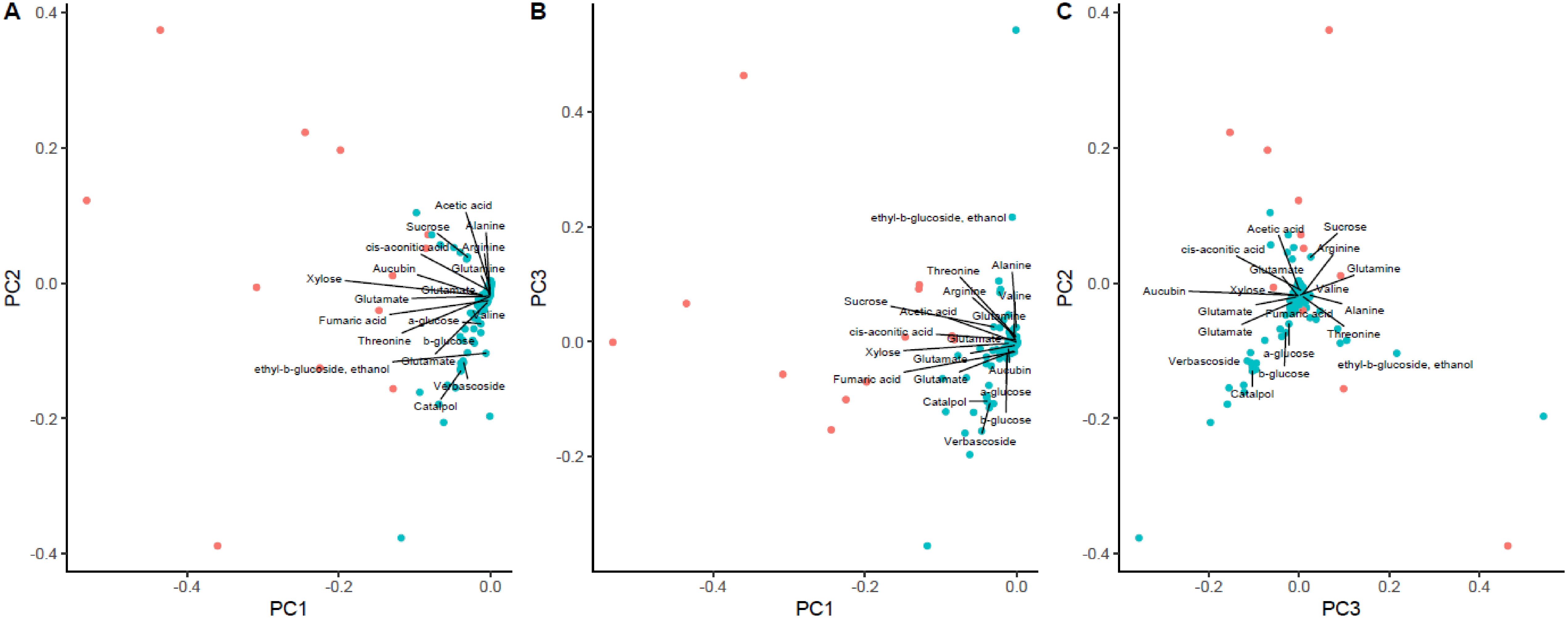
A
B
C
PC1
PC2
PC3
Acetic acid
Sucrose
Alanine
cis-aconitic acid
Arginine
Aucubin
Glutamine
Xylose
Glutamate
Valine
Fumaric acid
a-glucose
Threonine
b-glucose
ethyl-b-glucoside, ethanol
Verbascoside
Catalpol

Figure S3

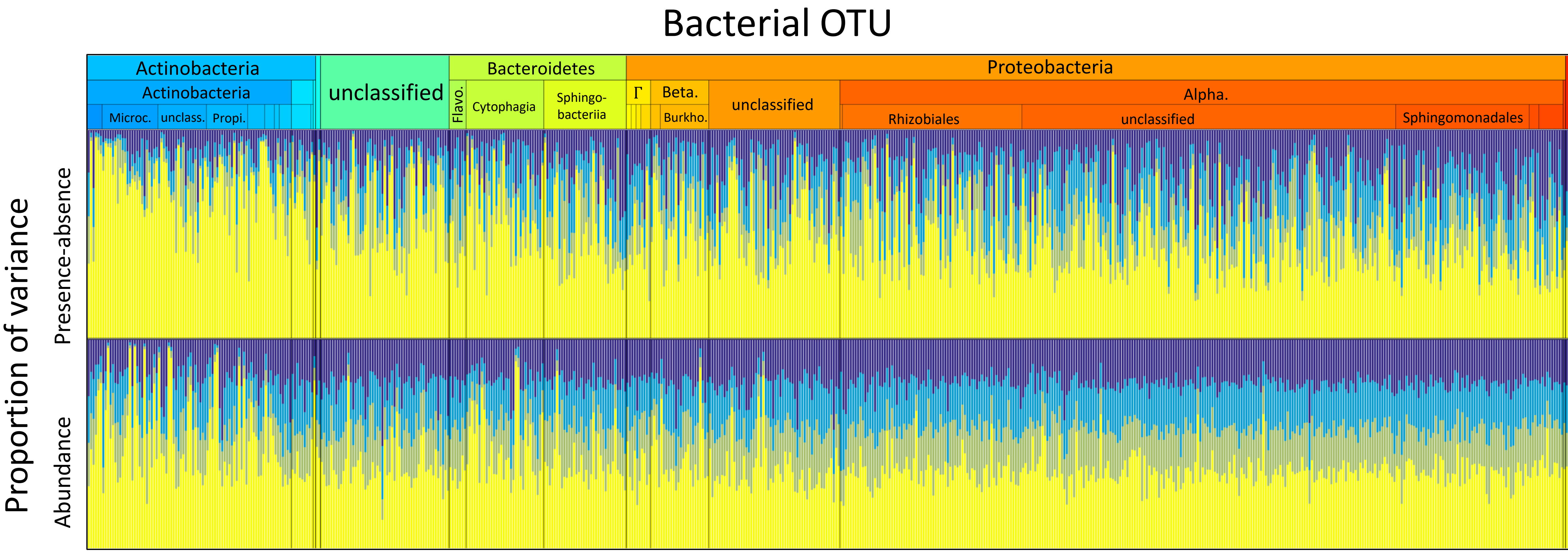


| Variable | Type | Color | P-A (%) | Ab (%) |
|---|---|---|---|---|
| Plant metabolic composition (PC1) | Fixed effect | | 18 | 21 |
| Plant metabolic composition (PC2) | Fixed effect | | 17 | 20 |
| Plant metabolic composition (PC3) | Fixed effect | | 14 | 19 |
| Plant level | Random effect | | 51 | 40 |

Figure S4

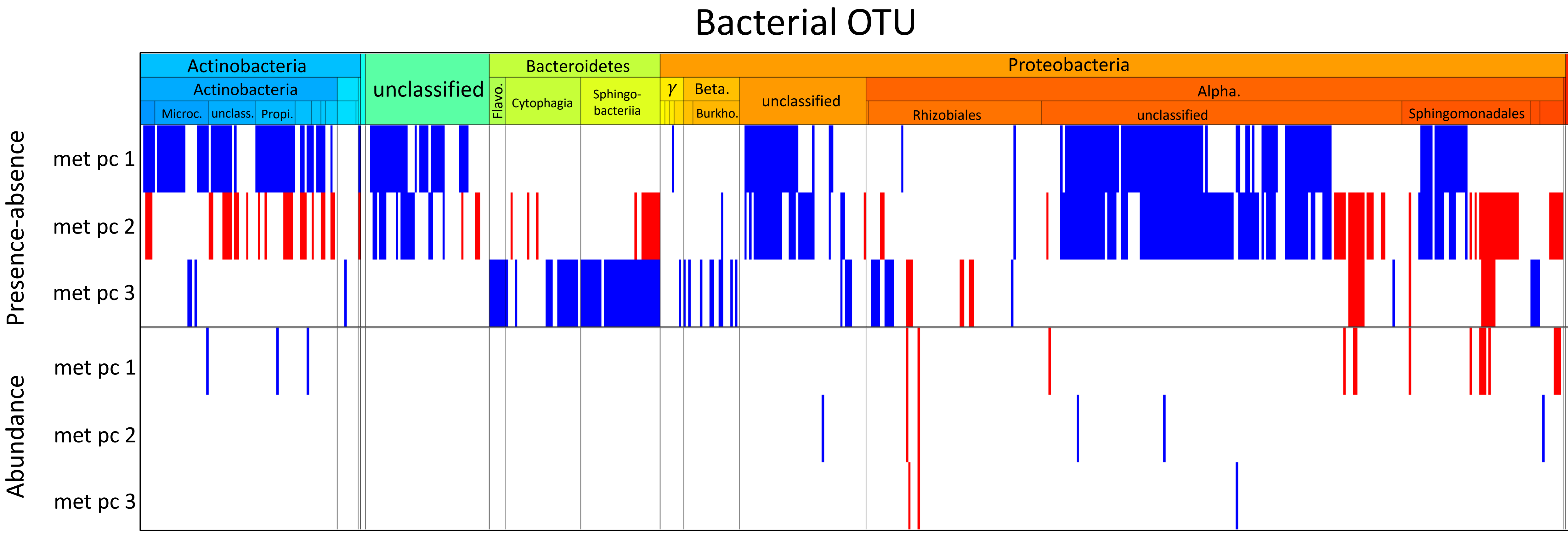
Bacterial OTU
Actinobacteria
Actinobacteria
Microc.
unclass.
Propi.
unclassified
Bacteroidetes
Flavo.
Cytophagia
Sphingo-
bacteriia
Proteobacteria
γ
Beta.
Burkho.
unclassified
Alpha.
Rhizobiales
unclassified
Sphingomonadales
Presence-absence
met pc 1
met pc 2
met pc 3
Abundance
met pc 1
met pc 2
met pc 3

Figure S5

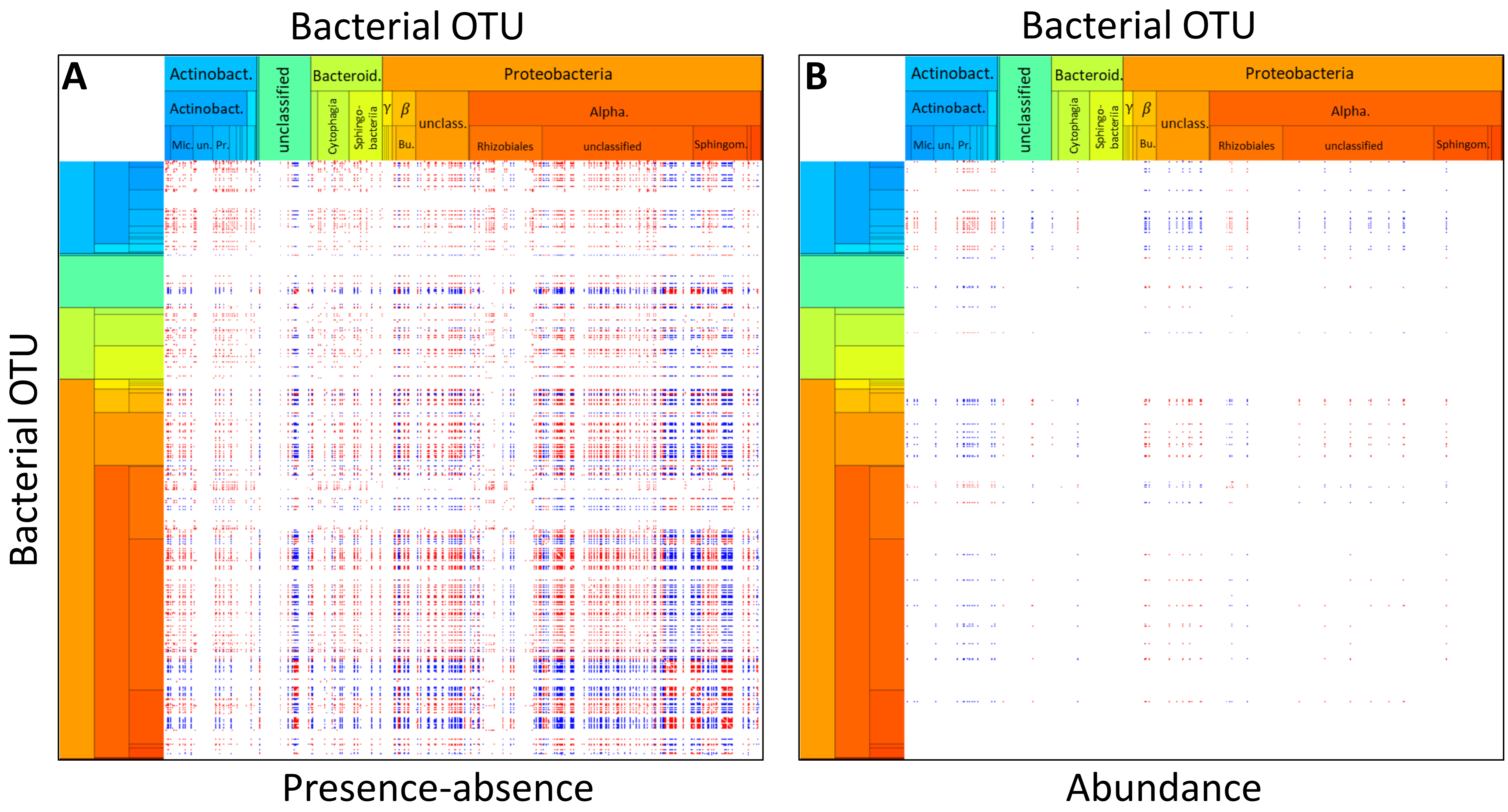
Bacterial OTU
A
Actinobact.
Actinobact.
Mic.
un.
Pr.
unclassified
Bacteroid.
Cytophagia
Sphingo-bacteriia
Proteobacteria
γ
β
Bu.
unclass.
Alpha.
Rhizobiales
unclassified
Sphingom.
Bacterial OTU
Presence-absence
Bacterial OTU
B
Actinobact.
Actinobact.
Mic.
un.
Pr.
unclassified
Bacteroid.
Cytophagia
Sphingo-bacteriia
Proteobacteria
γ
β
Bu.
unclass.
Alpha.
Rhizobiales
unclassified
Sphingom.
Abundance

Figure S6

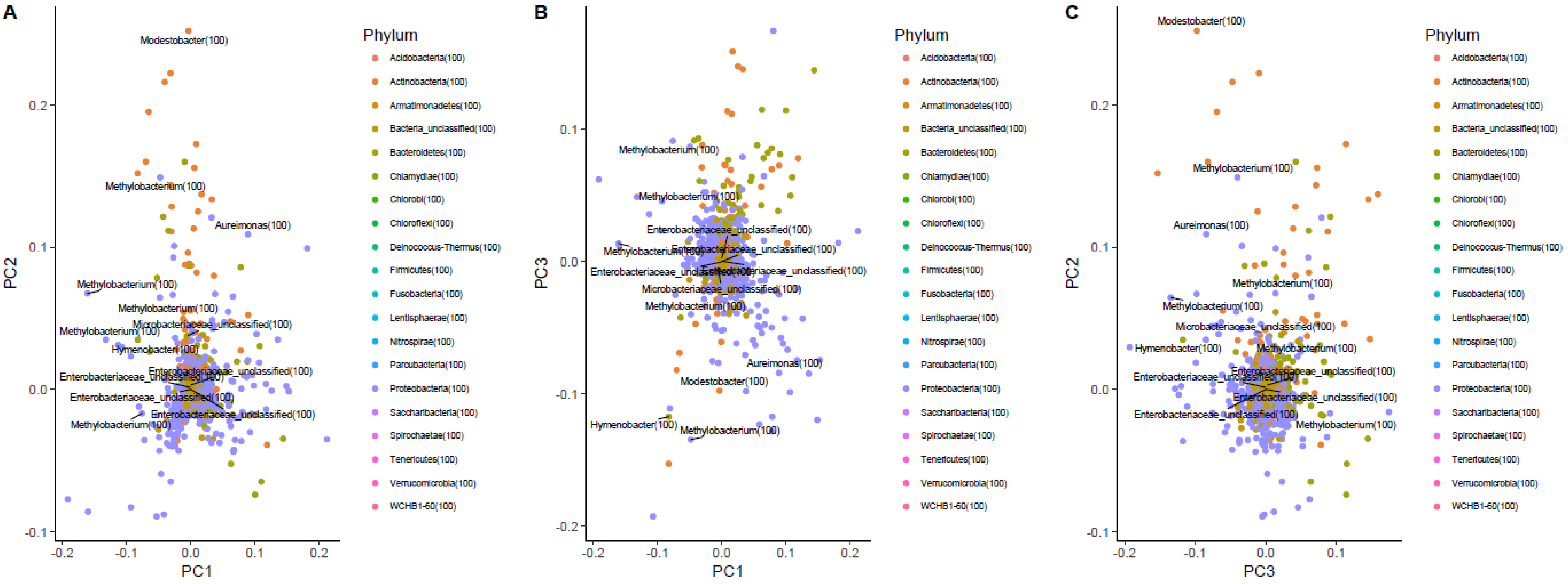

A
PC2
PC1
0.2
0.1
0.0
-0.1
-0.2
Modestobacter(100)
Methylobacterium(100)
Aureimonas(100)
Microbacteriaceae_unclassified(100)
Hymenobacter(100)
Enterobacteriaceae_unclassified(100)
B
PC3
PC1
C
PC2
PC3
Phylum
Acidobacteria(100)
Actinobacteria(100)
Armatimonadetes(100)
Bacteria_unclassified(100)
Bacteroidetes(100)
Chlamydiae(100)
Chlorobi(100)
Chloroflexi(100)
Deinococcus-Thermus(100)
Firmicutes(100)
Fusobacteria(100)
Lentisphaerae(100)
Nitrospirae(100)
Parcubacteria(100)
Proteobacteria(100)
Saccharibacteria(100)
Spirochaetae(100)
Tenericutes(100)
Verrucomicrobia(100)
WCHB1-60(100)


Figure S7

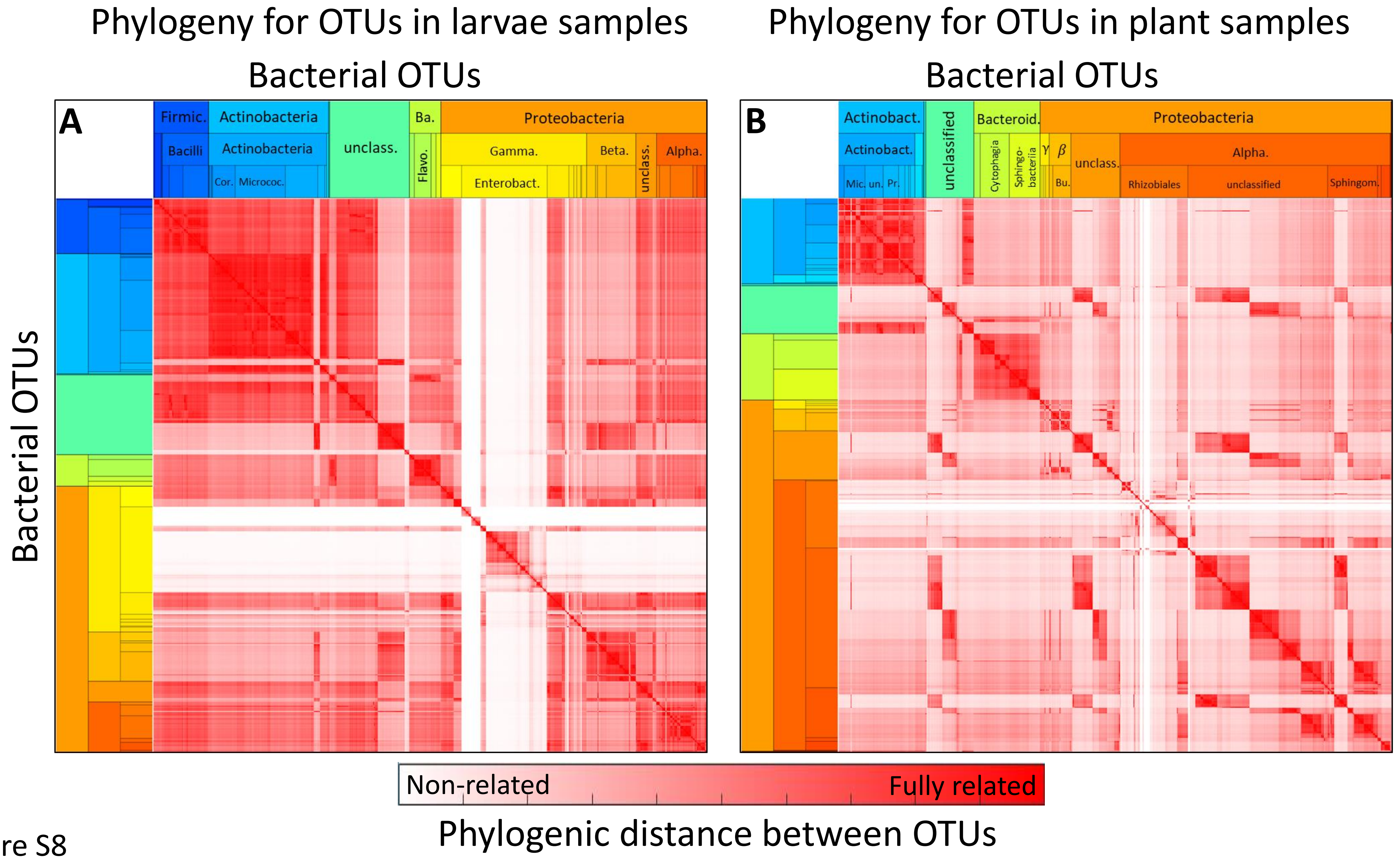
Phylogeny for OTUs in larvae samples
Bacterial OTUs
Phylogeny for OTUs in plant samples
Bacterial OTUs
A
Firmic.
Actinobacteria
unclass.
Ba.
Proteobacteria
Bacilli
Actinobacteria
Flavo.
Gamma.
Beta.
unclass.
Alpha.
Cor.
Micrococ.
Enterobact.
B
Actinobact.
unclassified
Bacteroid.
Proteobacteria
Actinobact.
Cytophagia
Sphingo-bacteriia
γ
β
unclass.
Alpha.
Mic.
un.
Pr.
Bu.
Rhizobiales
unclassified
Sphingom.
Bacterial OTUs
Non-related
Fully related
Phylogenic distance between OTUs

Figure S8
